# Dentate gyrus development requires a cortical hem-derived astrocytic scaffold

**DOI:** 10.1101/2020.06.11.145672

**Authors:** Alessia Caramello, Christophe Galichet, Karine Rizzoti, Robin Lovell-Badge

## Abstract

During embryonic development, radial glial cells give rise to neurons, then to astrocytes following the gliogenic switch. Timely regulation of the switch, operated by several transcription factors, is fundamental for allowing coordinated interactions between neurons and glia. We deleted the gene for one such factor, SOX9, early during mouse brain development and observed a significantly compromised dentate gyrus (DG). We dissected the origin of the defect, targeting embryonic *Sox9* deletion to either the DG neuronal progenitor domain or the adjacent cortical hem (CH). We identified in the latter previously uncharacterized ALDH1L1+ astrocytic progenitors, which form a fimbrial-specific glial scaffold necessary for neuronal progenitor migration towards the developing DG. Our results highlight an early crucial role of SOX9 for DG development through regulation of astroglial potential acquisition in the CH. Moreover, we illustrate how formation of a local network, amidst astrocytic and neuronal progenitors originating from adjacent domains, underlays brain morphogenesis.

## INTRODUCTION

Neuroepithelial cells (NECs) are the origin of all neurons, glia and stem cells found in the central nervous system (CNS)^1^. During early CNS development, NEC potential is initially restricted to a neuronal fate. But, at around E10.5 in the mouse, NECs undergo an irreversible switch, becoming radial glial cells (RGCs), which enable them to later generate astrocyte and oligodendrocyte progenitors^2^. Spatio-temporal control of the gliogenic switch is crucial because it regulates emergence and abundance of each cell type, enabling local establishment of fundamental neuron-glia interactions which are necessary for achieving correct CNS cytoarchitecture and functionality^3–5^. Amongst other roles, these interactions are essential to support the migration of differentiating neurons^6^. RGCs are known to guide migration of their neuronal progeny, while in the adult brain, astrocytes guide neuronal progenitors from the subventricular zone to the olfactory bulb^7, 8^. However, because astrocytic commitment has been difficult to monitor in the embryo, since specific markers to distinguish these from RGCs were lacking until recently^9, 10^, a role for astrocytes to support migration in the embryo had not been established^11^.

The dentate gyrus (DG) of the hippocampus, is a packed V-shaped layer of granule neurons involved in spatial memory formation and pattern separation^12^, which hosts a niche of radial glia-like cells supporting adult neurogenesis^13^. During embryonic development, DG granule neuron progenitors originate from the dentate neuroepithelium (DNE) or primary (1ry) matrix of the archicortex. From E14.5 in the mouse, pioneer intermediate progenitors (IPs), followed by neural stem cells (NSCs) delaminate to migrate extensively within the parenchyma, along the Dentate Migratory Stream (DMS) or 2ry matrix^14^. From E16.5, this mixed cell population, which also comprises post-mitotic neurons at this stage, ultimately reaches the brain midline forming the 3ry matrix, where they distribute in the upper then lower blades of the DG, and differentiate into PROX1+ granule neurons^15^. This process continues until after birth, when 1ry and 2ry matrices eventually disappear, and formation of new granule neurons will exclusively rely on local adult neurogenesis^16^. The cortical hem (CH), which is adjacent to the DNE and subsequently develops into the underlying fimbria, is a fundamental hippocampal organizer^17^. It gives rise to Cajal-Retzius (CR) cells, which regulate DG progenitor migration via secretion of Reelin and SDF1^18–20^. Progenitor migration, along the DMS, follows the track of a GFAP+ glial scaffold, which stretches from both the DNE and fimbrial epithelium towards and around the forming DG^21^. Although definitive proof is lacking, the shape and directionality of GFAP+ filaments within the scaffold suggest a supportive role for progenitor migration, both along the 2ry matrix and within the 3ry matrix^18, 19, 22–27^. Because GFAP is expressed by both RGCs and astrocytes, the cellular nature of the glial scaffold is unclear. However, deletion of the transcription factors NF1A/B, which regulate astrocyte gene expression^28^, partly affects formation of the scaffold, suggesting an astrocytic contribution^22, 23, 29^. The mechanisms explaining its formation also remain ambiguous, and it has been suggested to have a dual origin with a proximal fimbrial part deriving from the fimbrial glioepithelium, and a distal supragranular domain originating from DNE progenitors^19, 22, 27^. Therefore, clear evidence of both cellular identity and regional origin have been lacking.

SOX9, an SRY-related high mobility group (HMG) box (SOX) transcription factor, starts to be expressed in NECs around E9.5, just before their transition to RGCs and neuro-to-glia switch. We and others showed that SOX9 is required for this process, both within the embryonic brain^30, 31^ and spinal cord^28, 32^, because generation of astrocytes and oligodendrocytes is affected by its deletion. The role of SOX9 during early CNS development has been analyzed, in particular by conditional deletion using *Nestin-Cre*^33^. However, this Cre-driver only becomes active from E10.5, after the onset of SOX9 expression^30^. Consequently, the relatively mild effect of *Sox9* loss on astrogenesis in *Sox9^fl/fl^;Nestin-Cre* mutants^28, 32^ might also be explained by its early, albeit temporary expression.

To better understand the role of SOX9 in CNS development, including at early stages, we performed conditional deletion using *Sox1^Cre/+^*, which is active from E8.5 almost exclusively in the neural tube^34, 35^, and compared these with *Sox9^fl/fl^;Nestin-Cre* mutants. In contrast with the latter model^33^, all *Sox9^fl/fl^;Sox1^Cre/+^* mutant mice survived, allowing post-natal analysis. Reduced DG size was the most prominent defect in adult *Sox9^fl/fl^;Sox1^Cre/+^* mutant brains, but not in the few surviving *Sox9^fl/fl^;Nestin-Cre* animals, and this was already visible in newborns, suggesting an earlier developmental defect. While the emergence and differentiation of granule neuron progenitors were not affected in either *Sox9* mutant embryos, we observed that their migration within the developing DG was compromised, particularly in *Sox9^fl/fl^;Sox1^Cre/+^* mutants. We then showed that formation of the fimbrial glial scaffold, which is likely supporting neuronal progenitor migration towards the forming DG, was delayed in *Sox9^fl/fl^;Sox1^Cre/+^* mutants. We furthermore identified ALDH1L1+ astrocytic progenitors in the adjacent CH as the origin of the fimbrial glial scaffold. Accordingly, formation of these progenitors is significantly compromised in *Sox9^fl/fl^;Sox1^Cre/+^* mutants, but not in their *Sox9^fl/fl^;Nestin-Cre* counterparts, because *Nestin-Cre* is not active in the CH. Consequently, fimbrial glial scaffold and DG formation are less affected in these mutants. Exclusive deletion of *Sox9* in the CH, using *Wnt3a^iresCre^* ^17^, further confirmed that SOX9 is required for astrocyte progenitor emergence and hence fimbrial glial scaffold formation, allowing neuron progenitor migration. Ultimately, our results highlight the crucial importance of the timely emergence of glial progenitors in the CH for the establishment of a local supporting cellular network underlying neuronal migration during DG morphogenesis, and that, through its role in acquisition of astroglial potential, SOX9 is critical for this.

## RESULTS

### Adult DG morphology is sensitive to precise patterns of *Sox9* deletion in the archicortex

To further characterize the role of SOX9 during CNS development, we first performed an early CNS specific conditional deletion of the gene by crossing *Sox9^fl/fl^* ^36^ with *Sox1^Cre^*, which is active from E8.5^34^ prior to the onset of SOX9 expression^30^. The birth, growth and survival of *Sox9^fl/fl^;Sox1^Cre/+^* mice were not overtly affected. However, histological analyses of adult brains revealed that the DG was significantly reduced in size compared to controls (Fig. 1.Ai,ii;B). This was already evident at P2 (Fig. 1.Aiv-v, C), indicating this phenotype might arise earlier, due to the absence of SOX9 during DG embryonic development. Because the adult DG controls formation of new memories^12^, we assessed memory formation abilities in adult *Sox9^fl/fl^;Sox1^Cre/+^* mice by performing a Novel Object Recognition Test (NORT) (Fig. 1D). Failure to recognise a new object over a familiar one, detected as spending more time to investigate the new object, was demonstrated in *Sox9^fl/fl^;Sox1^Cre/+^* mice (Fig.1.E). Because deficiency in memory formation could also be caused by reduced exploration of the arena due to anxiety-like behaviours, we performed in parallel an open field test (Fig. S.1.A). This did not reveal any significant difference between control and mutant mice (Fig. S.1.B,C). Taken together these results show that embryonic deletion of *Sox9* affects memory forming abilities. This suggests that functionality of DG is affected, albeit we cannot exclude that defects in other regions of the mutant brains explain or exacerbate this behavioural deficiency.

**Figure 1.**
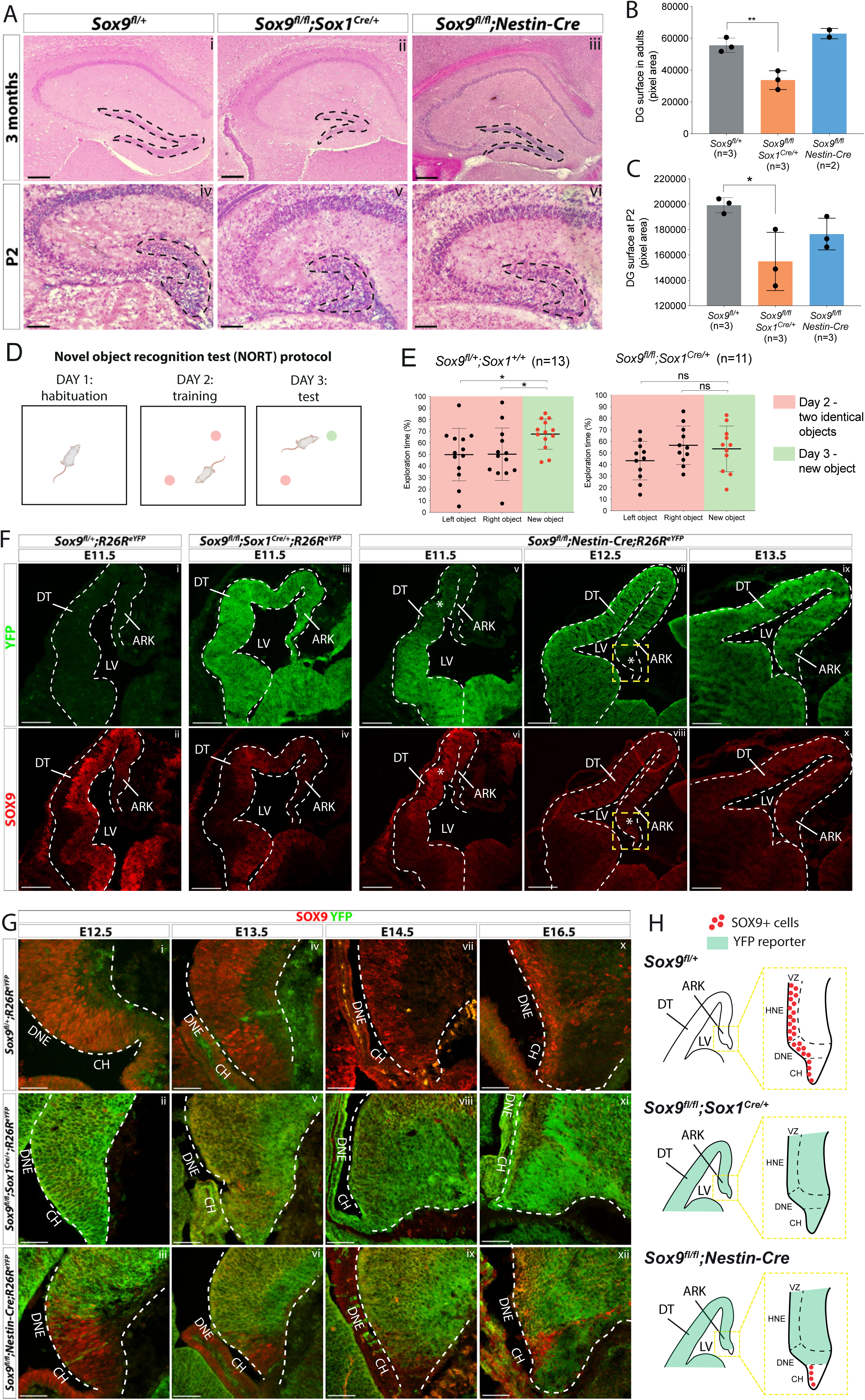
DG morphogenesis is differentially affected in *Sox1^Cre^* versus *Sox9^fl/fl^;Nestin-cre* mutants. **(A)** H&E staining of 3-month-old (i-iii) and P2 (iv-vi) brain sections of *Sox9^fl/fl^;Sox1^Cre/+^* and *Sox9^fl/fl^;Nestin-Cre* mutants compared to *Sox9^fl/+^* controls. DG (outlined) appears smaller in *Sox9^fl/fl^;Sox1^Cre/+^* mutants compared to both controls and *Sox9^fl/fl^;Nestin-Cre* mutants. **(B-C)** Quantification of DG surface as pixel area, in 3-month-old mice. (**B**) DG is significantly smaller in *Sox9^fl/fl^;Sox1^Cre/+^* mutants (33758±5898) compared to controls (55651±4492, P=0.0069, *t*-test) and *Sox9^fl/fl^;Nestin-Cre* mutants (62994±3243, statistical analysis for *Nestin-Cre* mutants is not possible as n<3). The defect is already visible in P2 pups (**C**), when DG area is significantly smaller in *Sox9^fl/fl^;Sox1^Cre/+^* mutants (154949±22878) compared to controls (199203±5990, P=0.029), but not compared to *Sox9^fl/fl^;Nestin-Cre* (176484±12459, P=0.277, Turkey’s multiple comparison test, ANOVA P=0.035). (**D**) Schematic of novel object recognition test (NORT) protocol. Pink and green circles represent familiar and new objects respectively. (**E**) Quantification of exploration times spent by mice over the identical left and right object on day 2 (red boxes) and on the new object on day 3 (green boxes). *Sox9^fl/+^;Sox1^+/+^* control mice (n=13) spend significantly more time exploring the new object on day 3 (67.55±13.16%) compared to time spend exploring the identical object on day 2 (left object: 49.83±22.65%, P = 0.0294; right object: 50.17±22.65%, P = 0.0391), indicating that they remember the objects from day 2. *Sox9^fl/fl^;Sox1^Cre/+^* mutants (n=11) instead do not remember the objects from day 2, because the time spent exploring the new object on day 3 (53.59±19.70%) is not different from the time spent exploring objects on day two, either on the left (43.37±16.65%, P = 0.2009) or right side (56.63±16.65%, P = 0.6839). **(F-H)** Immunofluorescence for YFP and SOX9 comparing respectively expression of *R26R^eYFP^* reporter of Cre activity and SOX9 expression patterns in *Sox9^fl/fl^;Sox1^Cre/+^*, *Sox9^fl/fl^;Nestin-Cre* mutants and controls, during forebrain (**F**) and archicortex (**G**) development. SOX9 remains expressed in *Nestin-Cre* mutants in both the CH and DNE (white asterisks in F). Yellow dashed square in (**F**) indicate area showed in (**G**) at higher magnification, also schematized in (**H**) together with *Sox1^Cre^* and *Nestin-Cre* recombination pattern in the ARK at E13.5. Signal from SOX9 immunofluorescence in *Sox9* mutant tissue was confirmed to be background with ISH for *Sox9* (Fig. S.3). LV: lateral ventricle; DT: dorsal telencephalon; ARK: archicortex; CH; cortical hem; DNE: dentate neuroepithelium; VZ: ventricular zone; HNE: hippocampal neuroepithelium; 1ry: primary matrix; 2ry: secondary matrix; 3ry: tertiary matrix. Scale bar represent 400µm in (Ai-iii); 200µm in (Aiv-vi) and (F); 50µm in (G).

*Sox9^fl/fl^;Nestin-Cre* mutants^30^ were generated in parallel to examine morphogenesis of the DG. In these, *Sox9* deletion occurs around 48 hours later than in *Sox9^fl/fl^;Sox1^Cre/+^* mutants, at E10.5, as SOX9 protein starts to be expressed in the developing CNS^30^. In contrast with *Sox9^fl/fl^;Sox1^Cre/+^* mutants, most *Sox9^fl/fl^;Nestin-Cre* animals die at birth^30^. Therefore, activity of *Nestin-Cre* outside the CNS^37^, in tissues where SOX9 is required, such as pancreatic islets^38^, heart^39^ and/or kidneys^40^, likely explains mortality in these mutants. We were, however, able to analyse two *Sox9^fl/fl^;Nestin-Cre* animals that survived until around 3 months of age and in which loss of SOX9 was confirmed by immunostaining (Fig. S.2). In contrast with *Sox1^Cre/+^* mutants, the size of the DG in *Sox9^fl/fl^;Nestin-Cre* animals was unaffected both in adults (Fig. 1.Aiii,B) and P2 pups (Fig. 1.Avi,C).

The difference in timing and/or pattern of embryonic *Sox9* deletion may underlie the variation in DG defects among these two mutant strains. To test this hypothesis, we analysed the activity of the *Sox1^Cre^* and *Nestin-Cre* by lineage tracing using an *R26^ReYFP^* allele and in parallel examined SOX9 expression. At E11.5, eYFP reporter expression was observed throughout the forebrain in *Sox9^fl/fl^;Sox1^Cre/+^;R26^ReYFP/+^* embryos (Fig. 1.Fiii) where, compared to controls, SOX9/*Sox9* expression was absent (Fig. 1.Fii,iv and Fig. S.3i-ii, B). However, in *Sox9^fl/fl^;Sox1^Cre/+^* mutant archicortices, we detected some rare *Sox9* positive cells by *in situ* hybridization (arrow in Fig. S.3.Aviii, see also Fig. 5.G), demonstrating that recombination of the *Sox9* allele is not quite ubiquitous. In contrast, *Nestin-Cre* is only active ventrally at E11.5, gradually progressing dorsally later in gestation (Fig. 1.Fv-x, Fig. S.3.Aiii,vi,ix). This implies that in *Sox9^fl/fl^;Nestin-Cre* mutants, SOX9 is transiently expressed in the DNE between E12.5 and E13.5 (Fig. 1.Giii,vi). In addition, the adjacent CH presents a mosaic pattern of recombination in *Nestin-Cre* mutants (Fig. 1.G.vi, ix, xii), as previously shown^19^. Consequently, many SOX9 expressing cells can still be found at E16.5 in this region in *Sox9^fl/fl^;Nestin-Cre* mutants (Fig. 1.Gxii).

Altogether these results suggest that delayed recombination in the DNE and/or residual expression of SOX9 in the CH of *Sox9^fl/fl^;Nestin-Cre* embryos (schematized in Fig. 1.H) underlies the difference in adult DG phenotype observed between *Sox9^fl/fl^*;*Sox1^Cre/+^* and *Sox9^fl/fl^;Nestin-Cre* mutants. Comparative analysis of DG development was then performed in both models to characterize the origin of the defect observed in *Sox9^fl/fl^*;*Sox1^Cre/+^* mutants.

### Abnormal distribution of granule neurons and their progenitors in the developing DG of *Sox9* mutants

DG progenitors, intermediate progenitors and differentiating granule neurons were first examined by analyzing respectively the expression of the transcription factors PAX6^41^, TBR2^42^ and PROX1^43^ at different stages of embryonic (E14.5, E16.5, E18.5) and post-natal (P2) DG development in *Sox9^fl/fl^;Sox1^Cre/+^* and *Sox9^fl/fl^;Nestin-Cre* mutants and controls (Fig. 2.A,F,I and Fig. S.4.A). All three cell types were found in the same numbers at embryonic stages in both *Sox9* mutants compared to controls (Fig. 2.E,G,J and Fig. S.4.B,C). However, at P2, TBR2+ intermediate progenitors are reduced in *Sox9^fl/fl^;Sox1^Cre/+^* mutants compared to controls, with 27% fewer cells (Fig. 2.G). Analysis of cleaved Caspase-3 immunostaining showed comparable patterns of cell apoptosis between mutants and controls, suggesting *Sox9* deletion does not affect progenitor survival at this stage (Fig. S.5). Moreover, Ki67 expression and EdU labelling, revealed no difference between controls and mutants in emergence, expansion and differentiation of DG granule neuron progenitors (Fig. S.6.A,B,D-F,H-J). Altogether, these data indicate while total number, survival and emergence of granule neurons and their progenitors are normal in both *Sox9^fl/fl^*;*Sox1^Cre/+^* and *Sox9^fl/fl^;Nestin-Cre* mutant embryos, there is a decrease in numbers of TBR2+ progenitors postnatally in *Sox9^fl/fl^;Sox1^Cre/+^* animals.

**Figure 2.**
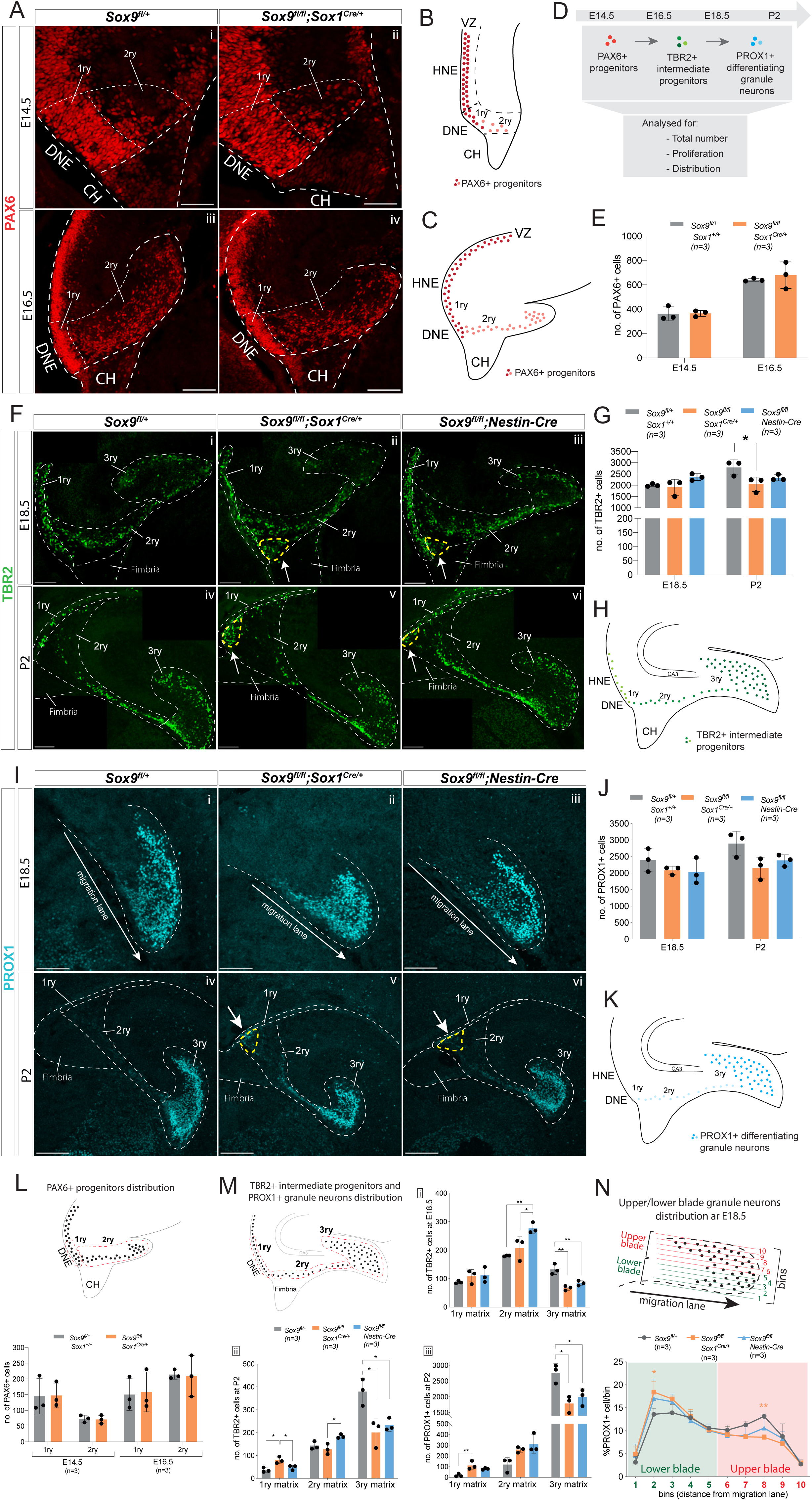
Granule neuron progenitors are generated normally but their distribution in SOX9 mutant developing DG is abnormal. **(A-K)** Immunofluorescence for PAX6 (**A**) TBR2 (**F**) and PROX1 (**I**) were performed at indicated developmental (E14.5: **A**i, ii; E16.5: **A**iii, iv; E18.5: **F**i-iii and Ii-iii) and postnatal stages (P2: Fiv-vi and Iiv-vi) of DG development. Arrows in **F**ii-iii,v-vi and Iv-vi point to respectively TBR2+ and PROX1+ cells accumulating close to the ventricle (yellow dashed line). 1ry, 2ry and 3ry matrices are delineated with white dashed lines. Localization within the developing DG of each cell type analyzed is schematized for PAX6 at E14.5 (**B**) and E16.5 (**C**), for TBR2 at E18.5 and P2 (**H**) and PROX1 for E18.5 and P2 (**K**), where color intensity in the illustration reflects level of markers expression. (**D**) shows experimental analysis. Quantification of total PAX6+ cells (**E**), TBR2+ cells (**G**) and PROX1+ cells (**J**) is shown at the indicated developmental and postnatal stages. Total cell number analysis shows a reduced number of TBR2+ cells at P2 (**G**) in *Sox9^fl/fl^;Sox1^Cre/+^* mutants (2041.67+323.72; P=0.03380 Dunn’s multiple comparison test; Kruskal-Wallis test P=0.0107) compared to controls (2792±331.72 TBR2+ cells). The same tendency was observed in P2 *Sox9^fl/fl^;Nestin-Cre* mutants (**G**) (2329.67±137.19 TBR2+ cells), and for PROX1+ cells (**J**) at P2 in both mutants *(Sox9^fl/fl^;Sox1^Cre^*: 2156.67±325.41 and *Sox9^fl/fl^;Nestin-Cre*: 2384.00 ± 173.25) compared to controls (2895.33±367.51). **(L, M)** Analysis of PAX6+, TBR2+ and PROX1+ cells distribution along the three matrices, according to the corresponding above schematics where dashed lines indicate areas considered for 1ry, 2ry and 3ry matrix quantification (also shown in **A**, **F**, **I**iv-vi). At E14.5 and E16.5 (**L**), the same amount of PAX6+ cells are found in the 1ry and 2ry matrix in *Sox9^fl/fl^;Sox1^Cre/+^* mutants compared to controls. At E18.5 (**M**i), more TBR2+ cells were found in the 2ry matrix of *Sox9^fl/fl^;Nestin-Cre* mutants (276.53±18.96) compared to both *Sox9^fl/fl^;Sox1^Cre/+^* mutants (207.33±39.85, P=0.03660) and controls (180.07±1.79, P=0.00850, Turkey’s multiple comparison test, ANOVA P=0.0090), while less TBR2+ cells were found in the 3ry matrix of both *Sox9^fl/fl^;Sox1^Cre/+^* mutants (66.93±7.90, P=0.0016) and *Sox9^fl/fl^;Nestin-Cre* mutants (84.00±8.50, P=0.0075) compared to controls (132.53±18.29, Turkey’s multiple comparison test, ANOVA P=0.0017). At P2 (**M**ii) more TBR2+ cells are found in 1ry matrix of *Sox9^fl/fl^;Sox1^Cre/+^* mutants (79.47+14.59), compared to controls (36.47±9.87, P=0.0101) and *Sox9^fl/fl^;Nestin-Cre* mutants (48.13±10.35, P= 0.0399, Turkey’s multiple comparison test, ANOVA P=0.0106). In *Sox9^fl/fl^;Nestin-Cre* mutants, more TBR2+ cells are accumulating in the 2ry matrix (184.07±8.47) compared to *Sox9^fl/fl^;Sox1^Cre/+^* mutants (127.87±22.72, P=0.0175, Turkey’s multiple comparison test, ANOVA P=0.0183). In both *Sox9* mutants, less TBR2+ cells are found in the 3ry matrix (*Sox9^fl/fl^;Sox1^Cre/+^*: 201.00±59.44, P=0.0119; *Sox9^fl/fl^;Nestin-Cre*: 233.73±27.81, P=0.029) compared to controls (378.93±57.88, Turkey’s multiple comparison test, ANOVA P=0.0109). At P2, PROX1+ cells (**M**iii) accumulate in the 1ry matrix of *Sox9^fl/fl^;Sox1^Cre/+^* mutants, (111.00±39.89) compared to controls (17.67±14.15, P=0.0088, Turkey’s multiple comparison test, ANOVA P=0.0100), and a significant decrease in the 3ry matrix of both *Sox9^fl/fl^;Sox1^Cre/+^* mutants (1786.67±266.25, P=0.0117) and *Sox9^fl/fl^;Nestin-Cre* mutants (1991.33±260.48, P=0.0329) is observed compared to controls (2758.33±297.16, Turkey’s multiple comparison test, ANOVA P=0.0112). **(N)** Analysis of the distribution of PROX1+ granule neurons distribution within the upper and lower blade of the forming DG at E18.5: the 3ry matrix was divided in 10 horizonal ventral to dorsal bins spanning the lower to upper blade domain. Cells were then counted within each bin. The percentage of PROX1+ cells present in each bin is represented. In *Sox9^fl/fl^;Sox1^Cre/+^* mutants, PROX1+ cells are accumulating in the lower blade (18.40±2.29%) compared to controls (13.57±1.29%, P=0.0187), and are reduced in the upper blade (8.57±0.58%) compared to controls (13.13±0.55%, P=0.0071, Turkey’s multiple comparison test, Two way ANOVA interaction P=0.0387, row factor P<0.0001, column factor P=0.9991). A similar tendency was observed in *Sox9^fl/fl^;Nestin-Cre* mutants, however it did not reached statistical significance. DG: dentate gyrus; DNE: dentate neuroepithelium; CH: cortical hem. Scale bar represent 100µm in (A), (F) and (Ii-iii); 200µm in (Iiv-vi).

While the total number of progenitors and granule neurons was unaffected in *Sox9* mutant embryos compared to controls, we observed an abnormal distribution of these cells along the three matrices (1ry, 2ry and 3ry) (Fig. 2.F,Iiv-vi,M). At E18.5, we counted more TBR2+ cells in the 2ry matrix in *Sox9^fl/fl^;Nestin-Cre* mutants, apparently at the detriment of the 3ry matrix, where fewer cells were present in both *Nestin-Cre* and *Sox9^fl/fl^;Sox1^Cre/+^* mutants compared to controls (Fig. 2.Mi). The misdistribution of TBR2+ cells become more evident post-natally, with fewer TBR2+ progenitors in the 3ry matrix of both mutant strains, but with more cells present in the 1ry matrix of *Sox9^fl/fl^;Sox1^Cre/+^* embryos and in the 2ry matrix of *Sox9^fl/fl^;Nestin-Cre* mutants, compared to controls (Fig. 2.Mii). We analyzed in parallel the distribution of TBR2+EdU+ progenitors at P2 and confirmed this was abnormal in *Sox9^fl/fl^;Sox1^Cre/+^* mutants, while not affected in *Sox9^fl/fl^;Nestin-Cre* mutants (Fig. S.6.G). Similarly, in both *Sox9* mutants at P2, we observed a reduction in PROX1+ differentiating granule neurons in the 3ry matrix and, in parallel, significantly increased number of these cells in the 1ry matrix of *Sox9^fl/fl^;Sox1^Cre/+^* mutants (Fig. 2.Miii). Conversely, 1ry to 2ry matrix distributions of PAX6+ and PAX6;Ki67+ progenitors (Fig. 2.L;Fig. S.6.C) are not affected in either E14.5 or E16.5 *Sox9^fl/fl^;Sox1^Cre/+^* mutants compared to controls. This suggests that defective DG neuronal progenitor distribution in *Sox9* mutants arise between E16.5 and E18.5.

Furthermore, the distribution of PROX1+ cells within the 3ry matrix also appears disrupted at E18.5 in both mutants, albeit less severely in *Sox9^fl/fl^;Nestin-Cre* embryos (Fig. 2.Ii-iii). To assess this defect, we quantified the number of PROX1+ cells in different bins ranging from the lower to the upper blade (Fig. 2.N). In controls, PROX1+ cells are equally distributed between the upper and lower blade. In contrast, in *Sox9^fl/fl^;Sox1^Cre/+^* mutants, these cells accumulate in the lower blade of the developing DG at the detriment of the upper blade. A similar tendency is observed in *Nestin-Cre* mutants; however, this does not reach statistical significance, suggesting a milder defect in this mutant strain. Furthermore, we noticed in the developing DG of both *Sox9* mutants, an ectopic cluster comprising a mix of both TBR2 and PROX1 expressing cells accumulating close to the ventricle since E18.5 (arrows in Fig. 2.F.ii,iii,v,vi, Iv,vi).

Altogether these analyses show that starting from E18.5 progenitors and granule neurons are abnormally distributed in the developing DG of *Sox9* mutants, with *Sox1^Cre^* mutants being more severely affected, both along the primary-to-tertiary matrix axis (Fig. 2.M) as well as within the forming DG (3ry matrix; Fig. 2.N). These observations suggest a defect in cell migration. The presence of an ectopic cluster of cells close to the ventricle is in agreement with this hypothesis, and we thus decided to characterize this further.

### Cell migration is impaired in the developing DG in absence of SOX9

We first analyzed the cellular composition of the ectopic cluster at P2 (Fig. 3.A-E). It is located next to the SOX2+ DNE (Fig. 3.Aiv-vi), and contains some SOX2+ progenitors and TBR2+ intermediate progenitors (some of which are EdU+; Fig. 3.A). Moreover, we also observed ectopic expression of PROX1 (arrows in Fig. 3.C) indicating that some progenitors are locally differentiating in granule neurons at this stage. Their commitment towards a granular cell fate was already visible at E18.5, with cells in the ectopic cluster expressing NeuroD1^44^ (arrows in Fig. 3.D). In conclusion, the ectopic cluster comprises cells at different stages of commitment toward the granule neuron fate.

**Figure 3.**
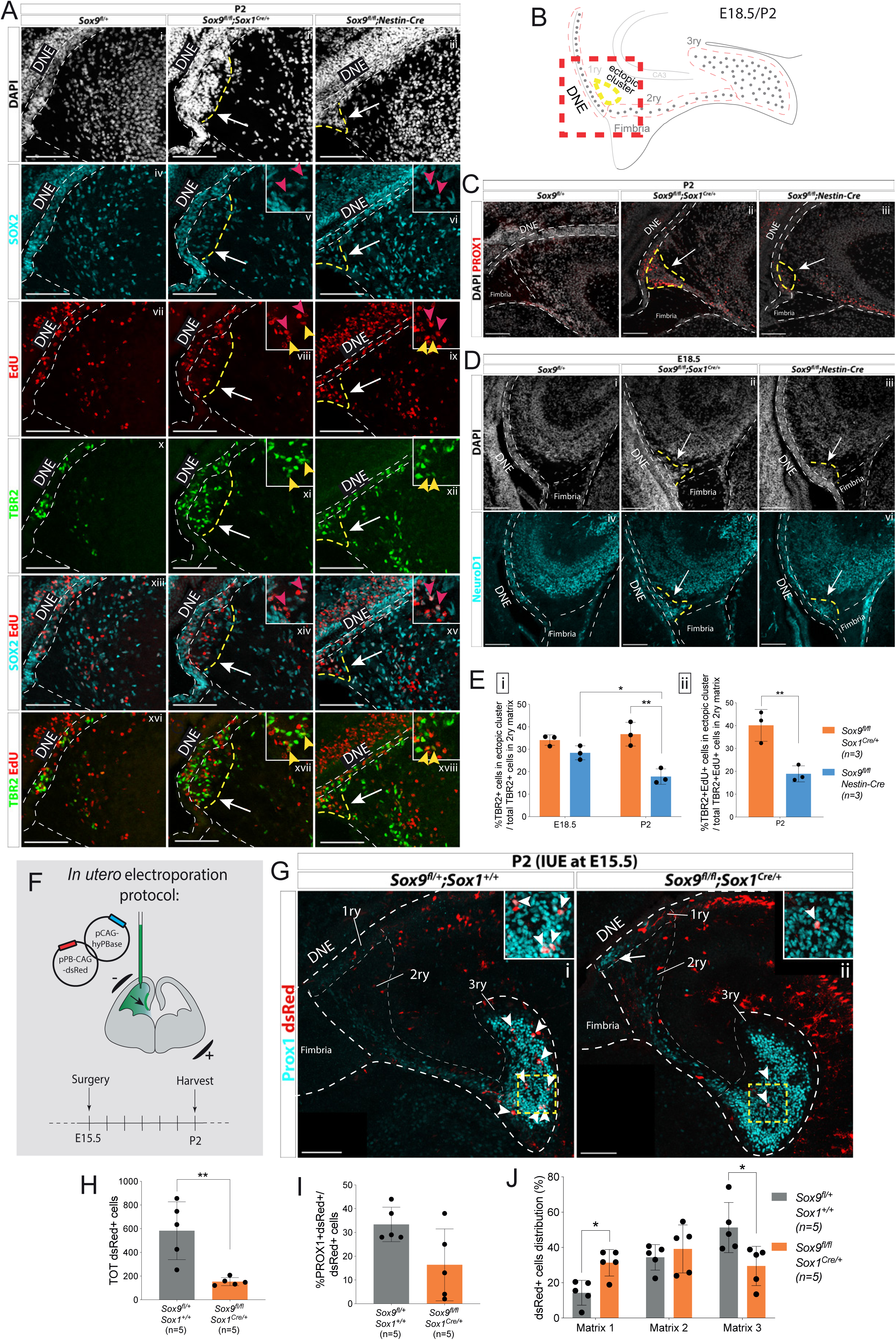
Ectopic accumulation of neuronal progenitors close to the ventricle suggests migratory defects in *Sox9* mutant DG. **(A)** Triple immunostaining for TBR2, SOX2 and EdU at P2 control, *Sox9^fl/fl^*;*Sox1*^Cre/+^ and *Sox9^fl/fl^*;*Nestin-Cre* brains. EdU was injected at E18.5. Insets show higher magnification of cells in ectopic cluster (schematized in **B**), magnified area is indicated by the white arrow. Yellow and pink arrowheads indicate respectively TBR2+EdU+ and SOX2+EdU+ cells in the ectopic cluster. **(B)** Illustration showing location of ectopic cluster within the developing DG. **(C-D)** Immunofluorescences for the differentiation markers PROX1 (**C**) and NeuroD1 (**D**) in *Sox9^fl/fl^;Sox1^Cre/+^* and *Sox9^fl/fl^;Nestin-Cre* mutants compared to controls at P2 (**C**) and E18.5 (**D**), respectively. Both markers are expressed by cells within the ectopic cluster (arrows) in *Sox9* mutants**. (E)** Quantification of ectopic cluster size at E18.5 and P2 in *Sox9^fl/fl^;Sox1^Cre/+^* compared to *Sox9^fl/fl^;Nestin-Cre* mutants. The percentage of TBR2+ progenitors in ectopic cluster relative to total number of TBR2+ progenitors in 2ry matrix is represented. At E18 (i), the ectopic cluster size was comparable between *Sox9^fl/fl^;Sox1^Cre/+^* (34.11±2.35%) and *Sox9^fl/fl^;Nestin-Cre* mutants (28.41±3.10%). It then significantly decreases in *Sox9^fl/fl^;Nestin-Cre* mutants at P2 compared to E18.5 (17.87±3.41%, P=0.0172, *t* test) and *Sox9^fl/fl^;Sox1^Cre/+^* at the same stage (36.73±5.30%, P=0.0061, *t* test). In agreement with the smaller ectopic matrix size at P2, less newly formed TBR2+EdU+ progenitors (ii) were found in the ectopic cluster of *Sox9^fl/fl^;Nestin-Cre* mutants (18.90±3.53%) compared to *Sox9^fl/fl^;Sox1^Cre/+^* (40.13±6.97%, P=0.0084 *t* test). **(F)** Schematic of *in utero* electroporation protocol. **(G)** Immunostaining for PROX1 and dsRed live fluorescence. Double positive cells from the dashed yellow square are showed at higher magnification in the inset. **(H)** The total number of dsRed+ cells was significantly smaller in *Sox9^fl/fl^;Sox1^Cre/+^* mutants (153.60±33.72) compared to controls (583.40±243.76, P=0.0045 *t* test). **(I)** The proportion of dsRed+ on total PROX1+ cells was not significantly reduced in *Sox9* mutants (16.40±15.11%) compared to controls (33.40±7.27%). **(J)** Distribution of dsRed+ cells along the three matrices (as schematized in Fig. 2.D). We observed more dsRed+ cells in the 1ry matrix of *Sox9^fl/fl^;Sox1^Cre/+^* mutants compared to controls (31.33±7.47% vs. 14.32±7.03%, P=0.0105 *t* test) and less in the 3ry matrix (29.49±11.13% vs. 51.27±14.20%, P=0.0287 *t* test). DNE: dentate neuroepithelium; IUE: *in utero* electroporation. Scale bars represent 100µm.

In our initial analysis of TBR2+ progenitor distribution (Fig. 2.M), cells present in the ectopic cluster were included into the 2ry matrix numbers (ectopic cluster + migrating cells within the 2ry matrix). To quantify the size of the ectopic cluster, which can be indicative of the migration defect, we calculated the percentage of TBR2+ cells clustering in the ectopic cluster relative to the total number of TBR2+ cells in the 2ry matrix. The size of the ectopic cluster represents a significant proportion of TBR2+ cells within the 2ry matrix, and was similar in both *Sox9^fl/fl^;Sox1^Cre/+^* (34.1±2.3%) and *Sox9^fl/fl^;Nestin-Cre* (28.4±3.1%) mutants at E18.5 (Fig. 3.Ei). While this proportion remained similar in *Sox9^fl/fl^;Sox1^Cre/+^* mutants at P2 (36.7±5.3%), it was significantly decreased in *Sox9^fl/fl^;Nestin-Cre* animals at the same stage (17.9±3.4%), in agreement with their milder phenotype (Fig. 3.Ei). A similar distribution was observed for newly generated progenitors at P2, with 40.13±6.79% of TBR2+EdU+ cells of the 2ry matrix clustering in the ectopic cluster in *Sox9^fl/fl^;Sox1^Cre/+^*mutants, while this proportion is more than halved in *Sox9^fl/fl^;Nestin-Cre* counterparts (18.0±3.5%; Fig. 3.Eii).

The presence of the ectopic cluster could be explained either by precocious differentiation of neuronal progenitors next to the ventricle or by their impaired migration toward the developing DG. To further analyze this aspect, lineage tracing of progenitors was performed using *in utero* electroporation in wild-type and *Sox9^fl/fl^;Sox1^Cre/+^* mutants. Progenitors facing the ventricle were electroporated at E15.5 with a plasmid expressing dsRed and the distribution of dsRed+ cells within the 3 matrices was analyzed 7 days later at P2 (Fig. 3.F,G). The total number of dsRed+ cells within the developing DG at P2 was significantly lower in *Sox9^fl/fl^;Sox1^Cre/+^* mutants compared to controls (Fig. 3.H). We did not observe excess cell death in electroporated mutants compared either to the contralateral side, or to controls at this stage (Fig. S.7). However, electroporated mutant progenitors, whose survival may have been compromised by lack of SOX9, may have been lost earlier. In controls, 33.4±7.26% of dsRed+ cells were PROX1+. In mutants, an average of 16.4±15.11% of dsRed+ cells were PROX1+, and this was not significantly different from controls. However, this proportion was variable. This may be explained by variability in the domain targeted by the electroporation since our previous data (Fig. 2.J) showed that loss of SOX9 does not have an effect on granule neuron differentiation. We then analyzed the distribution of dsRed+ cells in each matrix (represented by dotted white lines in Fig. 2.G). In P2 controls, the highest proportion of dsRed+ cells was observed in the 3ry matrix (51.27±14.20%), demonstrating that an important fraction of E15.5 electroporated progenitors had given rise to migrating granule neurons that successfully reached their destination in the developing DG (Fig. 3.G,J, arrowheads in G). In contrast, in *Sox9^fl/fl^;Sox1^Cre/+^* mutants, the highest proportion of dsRed+ cells was found in the 2ry matrix (39.18±13.59%) and the fraction of cells remaining in the 1ry matrix was significantly higher than in controls (Fig. 3.G,J). This suggests that, in *Sox9* mutants, a proportion of electroporated progenitors remained trapped near the DNE (arrow in Fig. 3.G.ii). These results are thus in agreement with impaired neuronal progenitor migration in the developing DG of *Sox9* mutants. We then investigated the origin of this phenotype by examining known molecular mechanisms regulating this process.

### Delayed induction of GFAP+ glial scaffold and its progenitors in absence of SOX9

Expression of chemokines (*Reln*, *Cxcl12*) and their receptors (*Vldlr*, *Cxcr4*) known to be involved in early stages of DG progenitor migration^26, 45^, is not significantly different in *Sox9* mutant E12.5 dissected archicortices compared to controls (Fig. S.8.A). These results are consistent with the absence of early migration defects in *Sox9* mutants (Fig. 2.L). Similarly, at E18.5, REELIN expression pattern and intensity appeared unchanged in both *Sox9* mutants compared to controls (Fig. S.8.B) further suggesting that CR cells are not affected by loss of *Sox9*.

In addition to CR cells, a GFAP expressing glial scaffold has been previously suggested to support DG progenitor migration during embryonic development ^22^. We thus examined expression of GFAP and observed a strongly positive scaffold in control samples from E18.5 connecting the DNE to the forming DG, through the fimbria (Fig. 4.Ai, B). In contrast, this is almost absent in *Sox9^fl/fl^;Sox1^Cre/+^* (Fig. 4.Aii) but only partially affected in *Sox9^fl/fl^;Nestin-Cre* embryos (quantified in Fig. 4.C), where the supragranular glial bundle is missing, but the fimbrial glial bundle is still visible (inset Fig. 4.Aiii; see schematic B). GFAP expression and scaffold structure in both *Sox9* mutants partially recover by P2 (Fig. 4.Aiv-vi, quantified in C), suggesting that absence of SOX9 might only delay scaffold formation, and that either compensatory or independent mechanisms may allow recovery early post-natally. However, the impact on DG morphogenesis is permanent.

**Figure 4.**
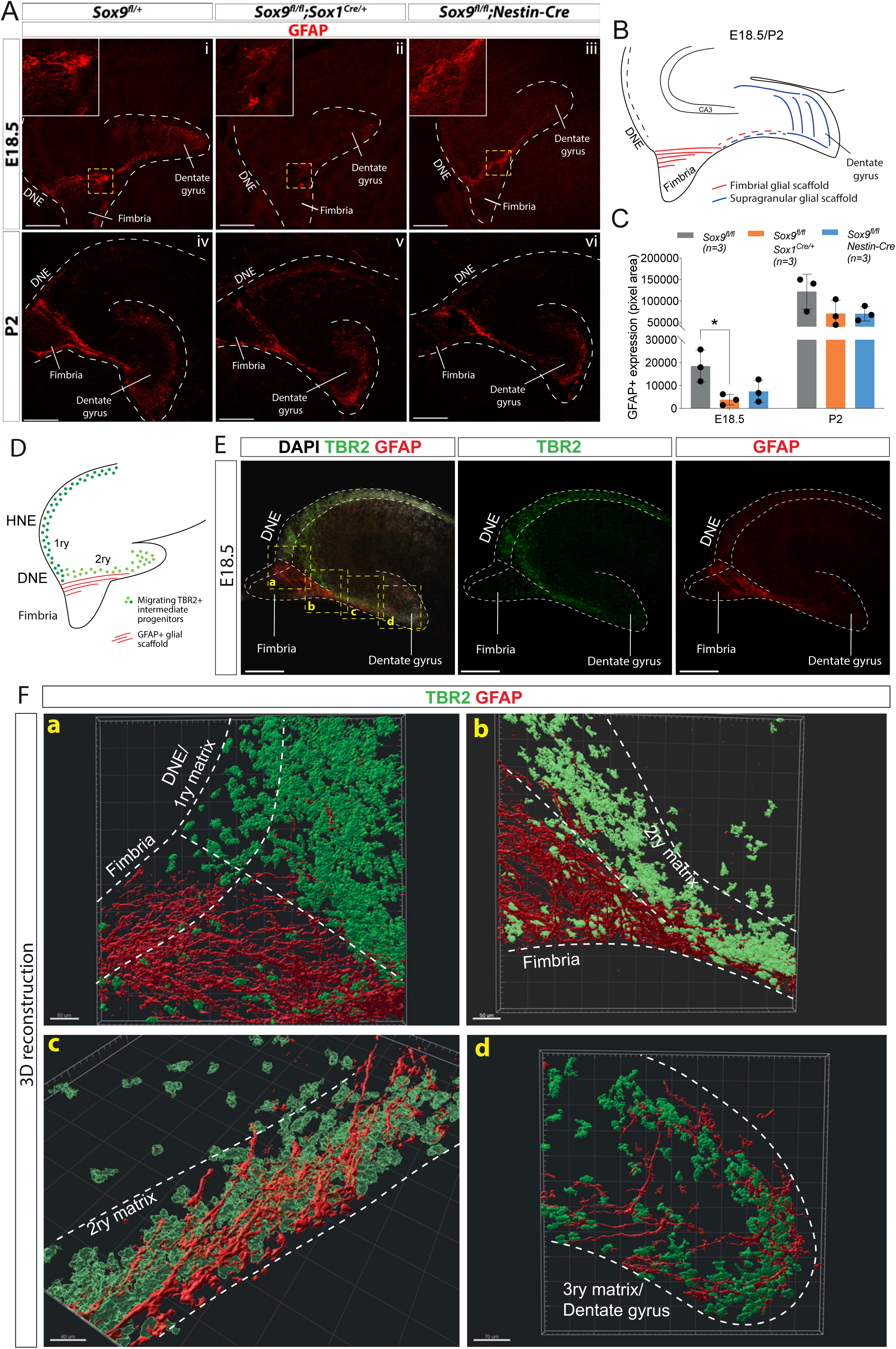
Delay in formation of the glial scaffold in *Sox9* mutants may explain progenitor migration defects. **(A-C)** Analysis of glial scaffold formation. **A)** Immunofluorescence for GFAP on E18.5 (**A**i-iii) and P2 (**A**iv-vi) control, *Sox9^fl/fl^;Sox1^Cre/+^* and *Sox9^fl/fl^;Nestin-Cre* brains showing GFAP reduction in both mutants at E18.5. Dashed line delineates the developing DG area, yellow dashed squares indicate areas magnified in insets. (**B**) Representation of the glial scaffold (red lines) in DG. (**C**) GFAP immunofluorescence quantification (pixel area). At E18.5, GFAP expression was significantly reduced in *Sox9^fl/fl^;Sox1^Cre^* mutants (18537.49±6990.47) compared to controls (3859.45±2347.25, P=0.0289), while quantification in *Sox9^fl/fl^;Nestin-Cre* did not reach statistical significance (7465.87±4875.50, P=0.0836, Turkey’s multiple comparison test, ANOVA P=0.0292). At P2, GFAP expression is recovered in both *Sox9* mutants compared to controls. (**D-F**) 3D reconstruction of control E18.5 embryos double immunostained for TBR2 and GFAP, (**E**) Representative control 10x single plane confocal images of sections processed for 3D reconstruction (schematized in **D**; yellow dashed squares indicate processed regions shown in **F**). (**F**) Snapshots from 3D reconstruction show that the fimbrial scaffold and 1ry matrix progenitors are initially separated (**a**). 2ry matrix migrating progenitors then start to intermingle with GFAP+ fibers as the scaffold elongates from the fimbria (**b**,**c**). 3ry matrix progenitors are also distributed within the supragranular scaffold within the developing DG (**d**). Movies of all 3D reconstructions are available in the supplementary material. DNE: dentate neuroepithelium. Scale bars represent 200µm in A and E.

The supporting role of the glial scaffold for DG neuronal progenitor migration has never been formally demonstrated. We aimed to further assess its functionality in this context by closely examining the distribution of TBR2 progenitors in relation to the GFAP+ scaffold at E18.5 when the scaffold first appears (Fig. 4.D-F). Close to the DNE, the GFAP+ fimbrial scaffold appears well separated from both the intermediate DNE progenitors and those migrating in the 2ry matrix (Fig. 4.Fa). Along the migratory stream in the 2ry matrix, progenitors start being more closely associated with the GFAP+ scaffold (Fig. 4.Fb). Finally, from the distal part of the fimbria, progenitors and scaffold appear completely intermingled (Fig. 4.Fc). We observe a similar association in the 3ry matrix between intermediate progenitors and the GFAP supragranular scaffold (Fig. 4.Fd). The position of intermediate progenitors relative to the glial scaffold suggest close contacts between these cell populations, supporting a functional role for the glial scaffold in promoting progenitor migration, particularly within the 2ry matrix. Consequently, delayed formation of the glial scaffold in *Sox9* mutants is likely implicated in the defective migration of DG progenitor.

Because SOX9 directly regulates the expression of *Gfap* in the developing spinal cord^28^, absence of GFAP expression in *Sox9* mutants may simply reflect downregulation of the gene. Therefore, to confirm the transient defect in glial scaffold formation, we examined the expression of ALDH1L1, an astrocyte specific marker^46^. At E18.5, ALDH1L1 expression pattern overlaps with that of GFAP, particularly within the fimbria where we see many GFAP+;ALDH1L1+ fibers (arrowheads in inset b in Fig. 5.Ai). In contrast, we observe GFAP+ALDH1L1-fibers around the developing DG (arrowheads in inset a in Fig. 5.Ai). At E16.5, before upregulation of GFAP, the ALDH1L1 expression pattern is reminiscent of that at E18.5, as mostly confined to the fimbria (Fig. 5.Aii). ALDH1L1+ cells are found as early as E13.5 in the archicortex (Fig. 5.Aiii) and also at this stage, they are specifically localized in the LEF1-negative;SOX2^high^ CH^47^ (Fig. 5.B,Ci-ii). The astrocytic nature of ALDH1L1+ cells was further confirmed by stainings for BLBP and GLAST, known markers of astrocytic progenitors^48^ which are also present within the LEF1-negative; SOX2^high^ CH (Fig. 5.Ciii,iv). Altogether, these results indicate that astrocytic progenitors are confined to the CH/fimbria throughout development suggesting they might later give rise to the fimbrial glial bundle.

**Figure 5.**
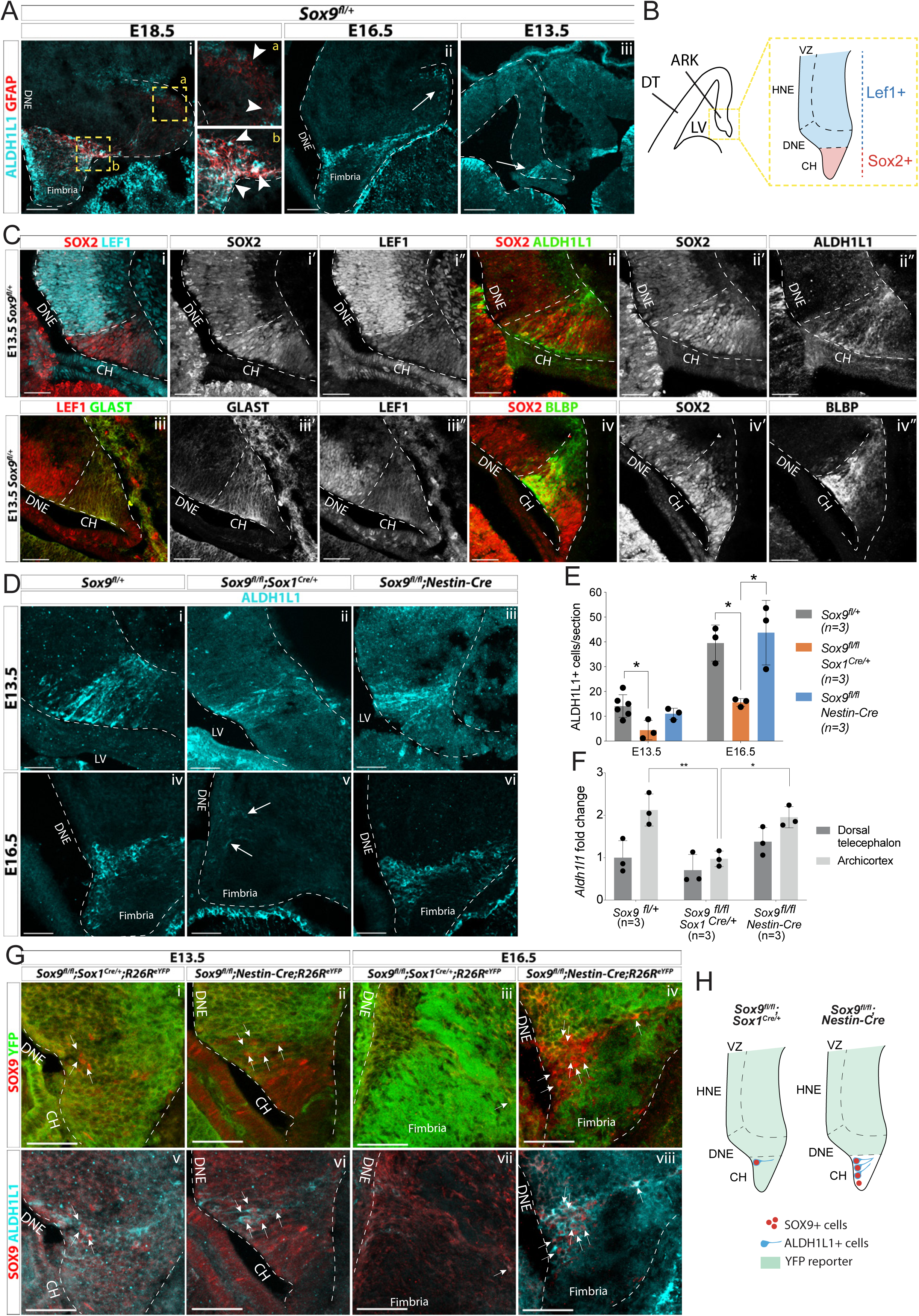
Emergence of astrocytic progenitor in the CH is affected in *Sox9* mutants according to levels of Cre activity in this region. **(A)** Double immunostaining for ALDH1L1 and GFAP in *Sox9^fl/+^* embryos at E18.5 (i), E16.5 (ii) and E13.5 (iii). ALDH1L1 and GFAP are co-expressed at E18.5 (i) in the fimbria (**b** insets on the right) but not around the forming DG (**a** insets on the right). Earlier, at E16.5 (ii), ALDH1L1, but not GFAP, is expressed in a similar pattern, in the fimbria and in a few cells around the forming DG (arrow in **A**ii), and as early as E13.5 in the archicortex (arrow in **A**iii). **(B, C)** Double immunostainings for SOX2;LEF1 (***i***), SOX2;ALDH1L1 (***ii***) at E13.5. SOX2 and LEF1 mutually exclusive expression patterns delineate the LEF1-SOX2^high^ CH and LEF1+SOX2^low^ DNE (schematized in **B**). ALDH1L1+ cells are exclusively located in the SOX2^high^ CH. Double immunostaining for GLAST;LEF1 (iii) and BLBP;SOX2 (iv) show a similar pattern of expression of the two astrocytic markers GLAST and BLBP in the LEF1-;SOX2^low^ CH, further suggesting ALDH1L1+ cells astrocytic nature. **(D-E)** Immunostaining (**D**) and quantification (**F**) of ALDH1L1+ cells in *Sox9* mutants at E13.5 (i-iii) and E16.5 (iv-vi) compared to controls. White arrows in **F**v indicate rare ALDH1L1+ cells found in *Sox9^fl/fl^;Sox1^Cre^* mutants. The number of ALDH1L1+ cells was significantly reduced in *Sox9^fl/fl^;Sox1^Cre^* mutants compared to controls, both at E13.5 (4.43±3.93 vs. 14.14±4.58, P=0.0193) and E16.5 (15.58±1.62 vs. 39.54±7.27, P=0.0338), while it was unaffected in *Sox9^fl/fl^;Nestin-Cre* mutants (E13.5: 11.00±2.29, P=0.5373, E16.5: 43.73±13.00, P=0.8288, Turkey’s multiple comparison test, ANOVA P=0.0242 and P=0.0147). **(F)** Analysis of *Aldh1l1* expression levels by qPCR from dissected DT and ARK of E12.5 *Sox9^fl/fl^;Sox1^Cre^, Sox9^fl/fl^;Nestin-Cre* and control embryos. *Aldh1l1* expression was significantly reduced in the ARK of *Sox9^fl/fl^;Sox1^Cre^* compared to both controls (P=0.0028) and *Sox9^fl/fl^;Nestin-Cre* mutants (P=0.0047, Turkey’s multiple comparison test, ANOVA P=0.0021). **(G,H)** Triple immunostaining for YFP, SOX9 and ALDH1L1 at E13.5 (i,ii,v,vi) and E16.5 (ii,iv,vii,viii) in *Sox9^fl/fl^;Sox1^Cre^* and *Sox9^fl/fl^;Nestin-Cre* mutants. A few double positive SOX9;ALDH1L1 cells are detected in the CH of both *Sox9* mutants (white arrows, schematized in **H**). More of these are present in *Sox9^fl/fl^;Nestin-Cre* compared to *Sox9^fl/fl^;Sox1^Cre/+^* mutants due differential Cre activity as shown by the *R26^ReYFP^* reporter expression. LV: lateral ventricle; DNE: dentate neuroepithelium; CH: cortical hem; DT: dorsal telencephalon; ARK: archicortex; HNE: hippocampal neuroepithelium; VZ: ventricular zone. Scale bars represent 200µm in (Aiii), 100µm in (Ai,ii) and 50µm in (C), (D) and (G).

We then analyzed whether ALDH1L1+ cells were affected by absence of SOX9. Strikingly, there was a dramatic reduction in their number in *Sox9^fl/fl^;Sox1^Cre/+^* mutants compared to controls, both at E13.5 and E16.5 (Fig. 5.D,E). Accordingly, *Aldh1l1* expression was significantly reduced in dissected archicortices of *Sox9^fl/fl^;Sox1^Cre/+^*

E12.5 embryos compared to controls (Fig. 5.F). This is consistent with a requirement for SOX9 for the emergence of astrocytic ALDH1L1+ progenitors, and consequently formation of the GFAP+ glial scaffold.

In contrast to *Sox9^fl/fl^;Sox1^Cre/+^* mutants, ALDH1L1+ cell number was unaffected in *Sox9^fl/fl^;Nestin-Cre* (Fig. 5.D-F). Because *Nestin-Cre*-mediated recombination occurs within the CH in a salt and pepper manner, SOX9 and ALDH1L1 expression patterns were analyzed in this region in both *Sox9* mutants. At E13.5, ALDH1L1+ cells were expressing SOX9 in the CH of *Sox9^fl/fl^;Nestin-Cre* embryos (Fig. 5.Gii,vi), and this was still observed at E16.5 (Fig. 5.Giv,viii). Interestingly, some rare SOX9+ cells were also present in the CH of E13.5 *Sox9^fl/fl^;Sox1^Cre/+^* mutants, and some were ALDH1L1+ (Fig. 5.Gi,v). These results suggest that in the CH of *Sox9* mutants, ALDH1L1+ cells may only arise from SOX9+ progenitors that escaped Cre recombination, which are present in higher numbers in *Sox9^fl/fl^;Nestin-Cre* compared to *Sox9^fl/fl^;Sox1^Cre/+^* mutants (schematized in Fig. 5.H). Moreover, the correlation between the extent of ALDH1L1+ cells and fimbrial glial scaffold loss with the severity of the progenitor migration defect in *Sox9^fl/fl^;Sox1^Cre/+^* versus *Sox9^fl/fl^;Nestin-Cre* mutants, further suggests a supporting migratory role of the scaffold.

Finally, in the developing spinal cord, the expression of the transcription factors NF1A/B are regulated by SOX9 and this is important for astrocytic differentiation^28^. We thus examined expression of NF1A and B in *Sox9^fl/fl^;Sox1^Cre/+^* E12.5 embryos. Both genes are expressed in the archicortex in control embryos. Loss of SOX9 does not affect either NF1A/B protein or transcript levels (Fig. S.9). We conclude that distinct molecular mechanisms downstream of SOX9 must underlie astrocytic specification in different domains of the CNS.

### CH-specific deletion of *Sox9* using *Wnt3a^iresCre/+^* impairs fimbrial glial bundle formation and compromises granule neuron progenitor migration

Because both *Sox1^Cre/+^* and *Nestin-Cre* are also active in the DNE (Fig. 1.G), cell autonomous defects could contribute to defective granule neuron progenitor migration. To examine this possibility and to confirm the requirement for SOX9 in the CH for formation of the fimbrial glial scaffold, CH-specific deletion of *Sox9* was performed using *Wnt3a^iresCre^* ^17^. First, we confirmed *Wnt3a^iresCre^* specificity to the CH by performing lineage tracing. In *Wnt3a^iresCre^*;*R26^ReYFP/+^* embryos, eYFP staining is mostly confined to the LEF1-CH at E12.5 (Fig. 6.Ai-iii) and to CH-derived REELIN+ Cajal-Retzius cells both around the DG (Fig. S.10.Ai-iv) and in the outer layer of the cortex (Fig. S.10.Av-viii). Because a few YFP+ cells were observed in the LEF1+ DNE, suggesting partial *Wnt3a^iresCre^* recombination in the DNE (arrowheads in Fig. 6.Ai-iii), we analyzed YFP expression in granule neurons at P2. Only 6.45±1% of PROX1+ granule neurons were YFP+ at this stage (arrowheads in Fig. 6.Bi-iii), suggesting *Wnt3a^iresCre^* is a suitable Cre driver for CH-specific deletion of *Sox9*.

**Figure 6.**
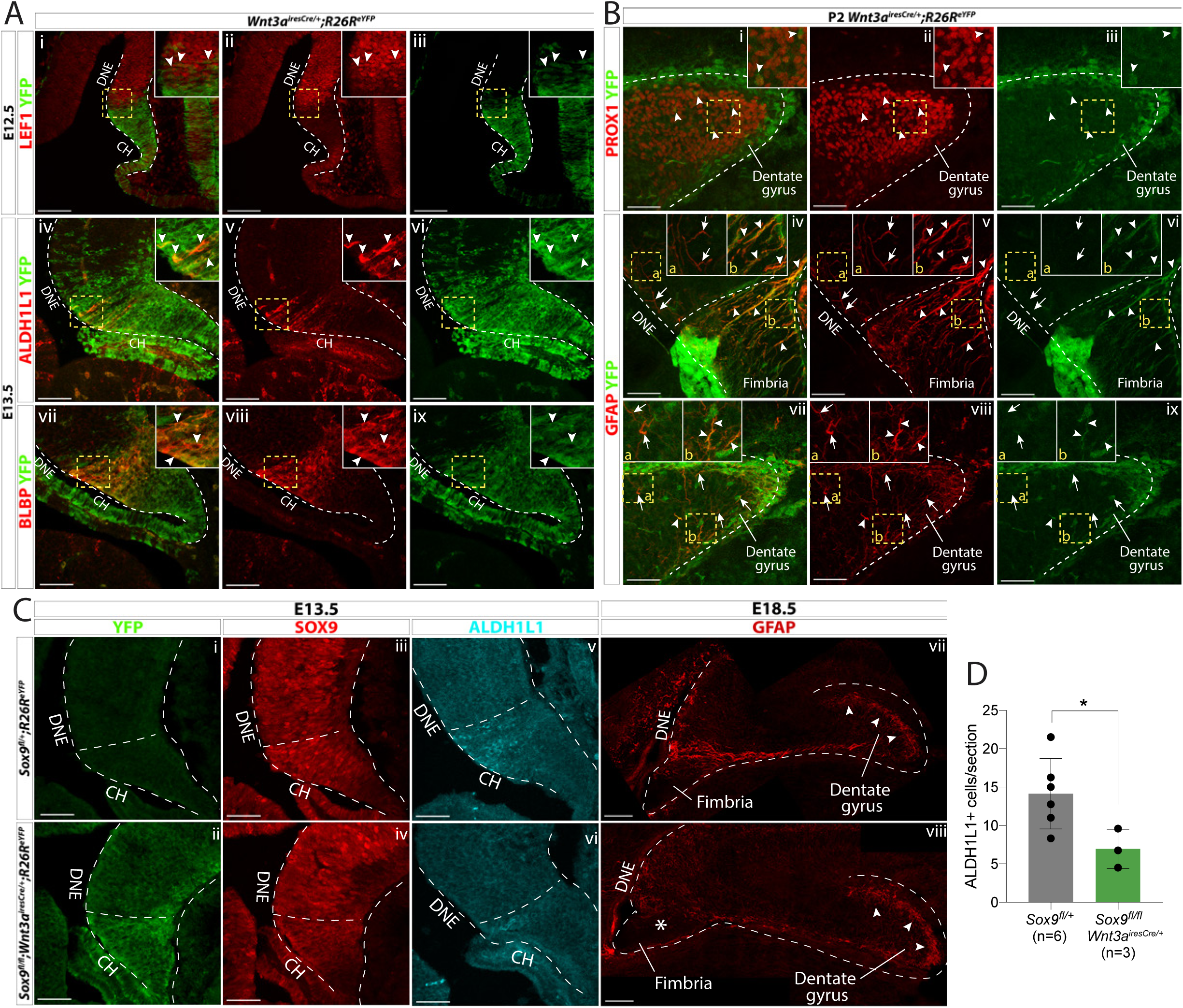
CH-specific deletion of *Sox9* using *Wnt3a^iresCre^* compromises glial scaffold formation exclusively within the fimbria. **(A,B)** Analysis of *Wnt3a^iresCre^* recombination pattern in the archicortex. (**A**) Double immunofluorescence for YFP with LEF1 (i-iii), ALDH1L1 (iv-vi) and BLBP (vii-ix) in *Wnt3a^iresCre/+^;R26R^eYFP^* embryos at E12.5 (i-iii) and E13.5 (iv-ix). Insets are magnified areas from yellow dashed boxes. Cre recombination is mostly observed in the LEF1-CH, however a few YFP+ cells are seen in the LEF1+ DNE (i-iii; arrowheads in magnified inset). ALDH1L1+;BLBP+ astrocytic progenitors express YFP in *Wnt3a^iresCre//+^;R26R^eYFP^* embryos (iv-xi; arrowheads in magnified inset), also confirming their CH origin. (**B**) Double immunofluorescences for PROX1;YFP (Bi-iii) and GFAP;YFP (iv,ix) in P2 *Wnt3a^iresCre/+^;R26R^eYFP^* embryos. Insets are magnified areas from yellow dashed boxes. Cells that have undergone Cre recombination are mostly GFAP+ and PROX1-, in agreement with a CH specific recombination pattern (**A**). Arrowheads in **B**i-iii and insets indicate some rare YFP+PROX1+ cells in the DG representing 6.45±1.00% of PROX1+ cells. Arrows and arrowheads in **B**iv,ix and insets indicates respectively YFP-GFAP+ and YFP+GFAP+ fibers in the DNE/fimbria (**B**iv-vi) and around the DG (**B**vii-ix), indicating the GFAP+ glial scaffold around the DG only partially originates from the CH. **(C,D)** Immunostainings and quantification for YFP (**C**i,ii); SOX9 (**C**iii,iv); ALDH1L1 (**C**v,vi) at E13.5 and GFAP at E18.5 in *Sox9^fl/fl^;Wnt3a^iresCre/+^;R26R^eYFP^* mutant compared to controls. In E13.5 archicortices of *Sox9^fl/fl^;Wnt3a^iresCre/+^* mutant and control embryos the CH specific deletion of SOX9 is confirmed. The number of ALDH1L1+ cells is significantly reduced (**D**) in *Sox9^fl/fl^;Wnt3a^iresCre/+^* mutants (6.95±2.56) compared to controls (14.14±4.58, P=0.022, *t*-test). At E18.5, the GFAP+ glial scaffold is affected exclusively within the fimbria (star in **C**viii) and not around the DG (arrowheads in **C**vii-viii). CH: cortical hem; DNE: dentate neuroepithelium. Scale bars represent 100µm in (Ai-and (Cvii-viii); 50µm in (Aiv-ix), (B), (Ci-vi).

ALDH1L1+ and BLBP+ astrocytic progenitors in the CH were also recombined by *Wnt3a^iresCre^* (Fig. 6.Aiv-ix). At P2, the fimbrial glial scaffold is entirely eYFP+ (arrowheads in Fig. 6.Biv-vi). We also observed some GFAP+;eYFP-filaments in the DNE that may represent DNE-derived radial glial cells (arrows in Fig. 6.Biv-vi). Conversely, the supragranular glial scaffold is made of both eYFP+ and eYFP-fibers suggesting a dual DNE and CH origin (respectively arrowheads and arrows in Fig. 6.Bvii-ix). Altogether these results suggest that the fimbrial glial scaffold is entirely derived from CH ALDH1L1+ astrocytic progenitors, while the supragranular glial scaffold only partially originates from CH. Additionally, we did not observe any PDGFRa+ oligodendrocyte precursor cells (OPCs) in the progeny of CH *Wnt3a^iresCre^* cells arguing in favor of an astrocytic, rather than radial-glial, nature of the scaffold.

*Sox9^fl/fl^;Wnt3a^iresCre/+^* mutants were then generated. While we were able to harvest mutant embryos until E18.5, animals died shortly after birth, precluding any post-natal analyses. *Wnt3a* is widely expressed in embryonic mesoderm precursors^49^ and deletion of *Sox9* in the embryonic heart, skeleton, pancreas and kidney is known to result in post-natal lethality^38–40^. In *Sox9^fl/fl^;Wnt3a^iresCre/+^* mutant CNS, SOX9 is absent specifically in the CH at E13.5 (Fig. 6.Ci-iv). Importantly, we also observe a 50% reduction of ALDH1L1+ cells in this area in E13.5 mutants compared to controls (Fig. 6.Cv,vi; quantified in D). Interestingly, at E18.5, the GFAP+ fimbrial glial scaffold is exclusively compromised in *Sox9^fl/fl^;Wnt3a^iresCre/+^* mutants (star in Fig. 6.Cviii), while the supragranular one is unaffected. These results are consistent with a role for SOX9 in the CH for specification of the ALDH1L1+ astrocytic progenitors giving rise to GFAP+ fimbrial glial scaffold. The normal appearance of the supragranular glial bundle in *Sox9^fl/fl^;Wnt3a^iresCre/+^* mutants is consistent with the observation that it may have a dual CH and DNE origin (Fig. 6.Bvii-ix).

We then analyzed granule neurons and their progenitors in *Sox9^fl/fl^;Wnt3a^iresCre/+^* mutants. At E18.5, the total number of PROX1+ granule neurons and TBR2+ progenitors is unchanged in mutants compared to controls (Fig. 7.A,B and Fig. 7.D,E respectively), similarly to what we observed in *Sox9^fl/fl^;Sox1^Cre/+^* mutants. Therefore, CH-specific deletion of *Sox9* does not affect progenitor formation and differentiation. We then analyzed the distribution of granule neurons. At E18.5, PROX1+ cell distribution in the 3ry matrix is unaffected in *Sox9^fl/fl^;Wnt3a^iresCre/+^* mutants compared to controls (Fig. 7.A,C). This is consistent with the supragranular glial scaffold not being affected upon CH-specific deletion of *Sox9* (Fig. 6.Fviii). In fact, repartition of granule neurons in the upper-blade and lower-blade at E18.5 is exclusively compromised in *Sox9^fl/fl^;Sox1^Cre/+^* and *Sox9^fl/fl^;Nestin-Cre* mutants (Fig. S.11). Because in both *Nestin*- and *Sox1^Cre^* models SOX9 is absent in the DNE, these results, together with previous observations^19, 27^, are in accord with a DNE contribution for the formation of the supragranular glial bundle.

**Figure 7.**
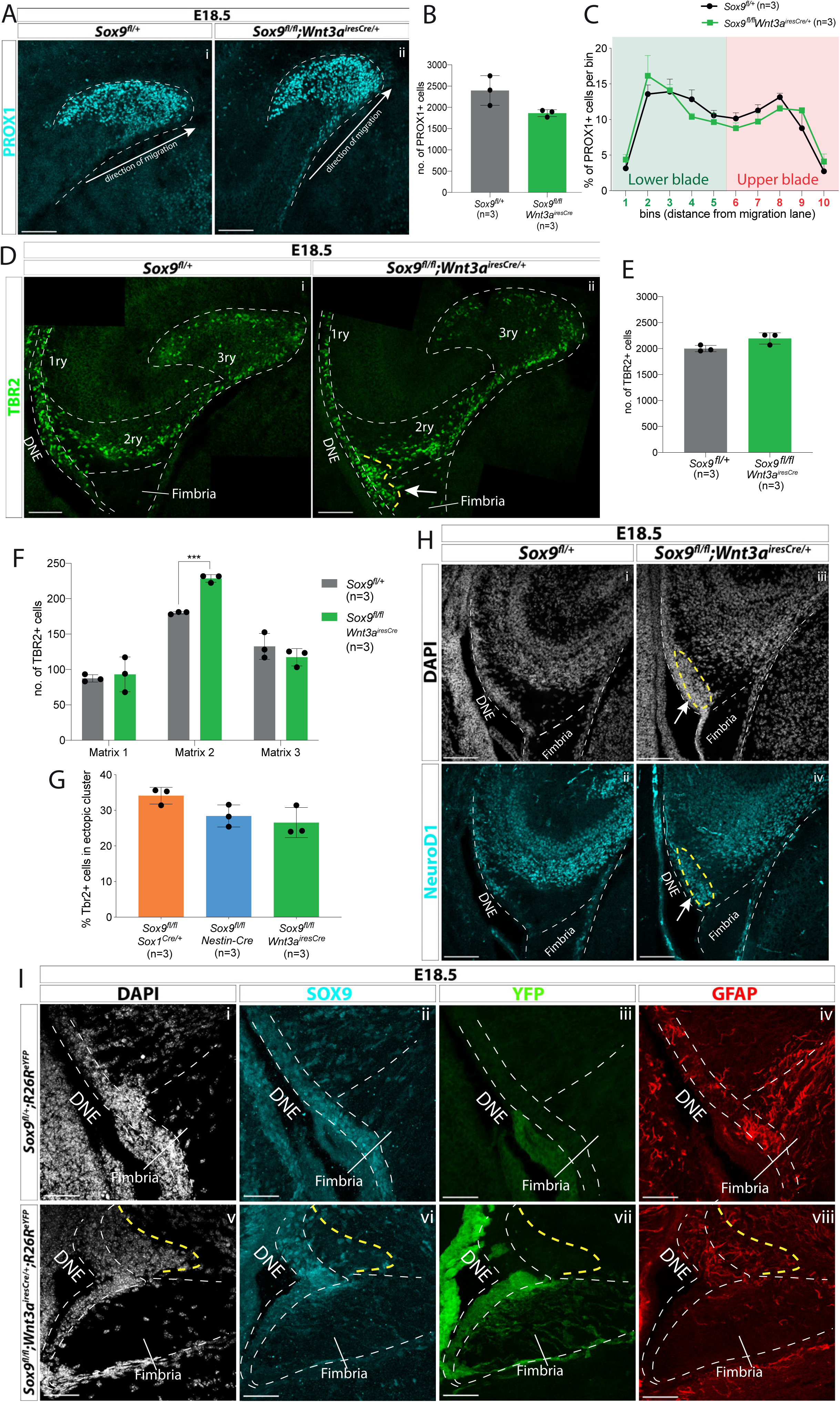
CH-specific deletion of *Sox9* using *Wnt3a^iresCre^* specifically affects granule neuron progenitor migration along the 1ry-to-3ry matrix axis. **(A-C)** Analysis of PROX1+ differentiating granule neurons in E18.5 *Sox9^fl/fl^;Wnt3a^iresCre/+^* DG. (**A**) Immunostaining for PROX1 on E18.5 controls and *Sox9^fl/fl^;Wnt3a^iresCre/+^* brains.. The total number of PROX1+ cells (**B**) and their distribution between upper and lower blade of the forming DG (**C**, see Fig. 3.E for analysis settings) was not affected in *Sox9^fl/fl^;Wnt3a^iresCre/+^* mutants compared to controls. (**D-G**) Analysis of TBR2+ intermediate progenitors at E18.5 in *Sox9^fl/fl^;Wnt3a^iresCre/+^* DG via immunofluorescence (**D**). The total number of TBR2+ cells is unchanged (**E**) but their distribution along the three matrices (**F**) is affected as there were more cells in the 2ry matrix of *Sox9^fl/fl^;Wnt3a^iresCre/+^* mutants (228.60±5.37) compared to controls (180.07±1.79, P=0.0001, *t* test). Arrow indicates accumulation of TBR2+ cells in the ectopic cluster in *Sox9^fl/fl^;Wnt3a^iresCre/+^* mutants. **G)** Percentage of TBR2+ in ectopic cluster. The percentage of TBR2+ cells in the ectopic cluster is comparable to that observed *Sox9^fl/fl^;Sox1^Cre/+^* and *Sox9^fl/fl^;Nestin-Cre* mutants (calculated as % of TBR2+ progenitors in ectopic cluster relative to total number of TBR2 progenitors in 2ry matrix). **(H)** Immunofluorescence for NeuroD1 showing ectopic differentiation towards granule neuron cell fate in the ectopic cluster of *Sox9^fl/fl^;Wnt3a^iresCre/+^* mutants (arrow). **(I)** Triple immunostaining for SOX9, YFP and GFAP on E18.5 controls and *Sox9^fl/fl^;Wnt3a^iresCre/+^* brains showing YFP-cells accumulating next to the SOX9+ DNE in E18.5 *Sox9^fl/fl^;Wnt3a^iresCre/+^;R26R^eYFP^* mutants (delineated by yellow dashed line) and underlaid by a defective GFAP scaffold. DNE: dentate neuroepithelium. Scale bars represent 50µm in (A), (Bi,ii) and (L); 100µm in (Biii,iv), (D), (G), and (J).

**Figure 8.**
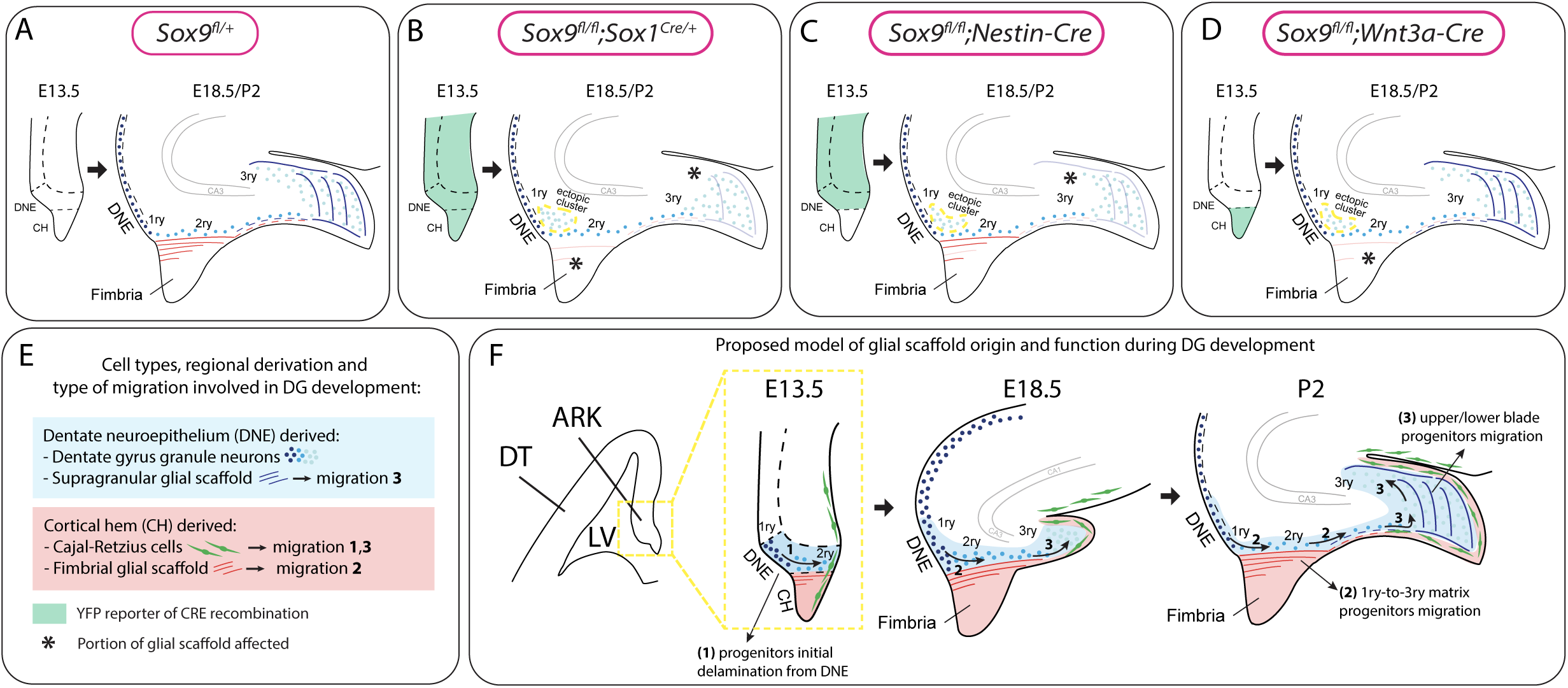
Model for the dual origin and function of the DG glial scaffold based on the analysis of defects following differential deletion of *Sox9*. **(A-D)** Schematic of mouse models used for *Sox9* conditional deletion and analysis of DG development. The pattern of Cre recombination is represented in the archicortex at E13.5 (green area) and the corresponding phenotype observed at E18.5/P2. Stars indicate local absence of the GFAP+ glial scaffold. **(E)** Figure legend. **(F)** Model of DG development based on defects observed following differential deletion of *Sox9*. At early stages of DG development (E13.5), granule neuron progenitors undergo delamination from the 1ry matrix and form the 2ry matrix (migration direction depicted by arrow **1**). This initial step is hypothesized to be independent of the glial scaffold because it is not affected in its absence in *Sox9* mutants. From E18.5, progenitor migration towards the forming DG/3ry matrix relies on the fimbrial scaffold (red lines, arrow **2**). This fimbrial scaffold derives from astrocytic progenitors located in the CH (red area). At the same time, the dentate scaffold around the DG (blue lines) provides support for granule neuron positioning within upper and lower blades of the forming DG (arrows **3**). Cells giving rise to this second scaffold are DNE derived (blue area). CH: cortical hem; DNE: dentate neuroepithelium; HNE: hippocampal neuroepithelium; VZ: ventricular zone; DT: dorsal telencephalon; ARK: archicortex.

We then examined distribution of TBR2+ progenitors in the three matrices at E18.5. We observe that more cells accumulate in the 2ry matrix of *Sox9^fl/fl^;Wnt3a^iresCre/+^* mutants compared to controls (Fig. 7.F). This abnormal distribution is reminiscent of that seen in *Sox9^fl/fl^;Sox1^Cre/+^* mutants (Fig. 2.F,M). Furthermore, progenitors in *Sox9^fl/fl^;Wnt3a^iresCre/+^* mutants form a cluster close to the ventricle (the ectopic cluster; Fig. 7.Dii), with a size comparable to that seen in both *Sox9^fl/fl^;Sox1^Cre/+^* and *Sox9^fl/fl^;Nestin-Cre* mutants at the same stage (Fig. 7.G). The ectopic cluster comprises differentiating neurons, with some cells expressing NeuroD1 (arrow in Fig.7.H) as previously observed in *Sox9^fl/fl^;Sox1^Cre/+^* and *Sox9^fl/fl^;Nestin-Cre* mutants (Fig. 3.D). Defective localization is not due to cell autonomous defects, because cells accumulating in the ectopic cluster are reporter negative in E18.5 *Sox9^fl/fl^;Wnt3a^iresCre/+^;R26R^eYFP^* mutants (Fig. 7.Iv-vii), indicating their precursors were not deleted for *Sox9*. Altogether, these results strongly suggest that SOX9 is required in the CH for astrocytic specification and subsequent fimbrial glial scaffold formation. Furthermore, they demonstrate that defective localization of neuronal progenitors is a non-cell-autonomous defect, consistent with a lack of migratory support by the defective CH-derived fimbrial glial scaffold (Fig. 7.Iviii).

## DISCUSSION

Using conditional deletion approaches, we have dissected the role of SOX9 during DG development. Differential patterns of gene deletion in the archicortex had marked consequences on adult DG morphology, and functionality, and we were consequently able to establish that SOX9 expression is required in the CH for proper DG morphogenesis. More precisely, our results highlight the crucial role of SOX9 for timely induction of the gliogenic switch in the CH, allowing emergence of astrocytic progenitors and subsequent formation of the fimbrial glial scaffold. Moreover, we show that this portion of the scaffold, which is closely associated with granule neuron progenitors, is necessary for their migration toward the forming DG. Furthermore, while partial recovery of the scaffold is observed early post-natally in *Sox9* mutants, DG morphogenesis are permanently affected, demonstrating that while the glial scaffold is only required transiently, it is indispensable. In conclusion, our study unravels the cascade of events orchestrating the establishment of supportive astrocytic-neural interactions required for DG morphogenesis, highlighting its sensitivity to timing and dependence on SOX9.

### Dual origin, function and nature of the DG glial scaffold

Lineage tracing experiments using *Wnt3a^iresCre^* demonstrate here that the fimbrial part of the glial scaffold has a CH origin. Our analyses of the scaffold and of the defects observed in its absence, show that it supports neuronal progenitor migration from 1ry-to-3ry matrix. This had been suggested previously, but without direct evidence^19, 22^. In addition, because in *Sox9* mutants, early PAX6+ progenitor migration is not affected and the ectopic cluster appears from E18.5, the fimbrial glial scaffold only become indispensable for migration at late stages of DG development. We hypothesize this may be explained by the increasing distance between DNE and the forming DG, as development proceeds.

In agreement with a DNE origin for the distal part of the scaffold^19, 27^, the supragranular glial bundle, was mostly unlabelled in our *Wnt3a^iresCre^* lineage tracing experiments. However, the presence of some labelled fibers suggests a CH contribution. Beside a different cellular composition, the supragranular scaffold has also been suggested to have a distinct function from the fimbrial one, where the former guides granule neuron migration within DG upper and lower blades^27^. In agreement, in *Sox9^fl/fl^;Wnt3a^iresCre/+^* mutants, where SOX9 is deleted exclusively in the CH, the supragranular scaffold and consequently distribution of granule neurons in the 3ry matrix of the developing DG, appear normal. In contrast, we observe a transient impairment of the supragranular scaffold in both *Sox9^fl/fl^;Sox1^Cre/+^* and, to a lesser extent, in *Sox9^fl/fl^;Nestin-Cre* mutants, along with an altered upper/lower blade distribution of granule neurons. These results are in accord with a different role for this part of the scaffold as well as its predominant DNE origin since both *Sox1* and *Nestin-Cre* drivers are active in this domain. However, we observe a milder alteration in DG upper/lower blade granule neuron distribution in *Sox9^fl/fl^;Nestin-Cre*. Because *Nestin-Cre* is not active in this domain, this result may be explained by a contribution of CH-derived SOX9+ glial cells to formation of the supragranular scaffold, in further agreement of a mixed DNE/CH origin for this distal scaffold.

ALDH1L1, GLAST, BLBP and GFAP co-expression in the fimbrial scaffold and its CH progenitors argue strongly for an astrocytic identity, demonstrating for the first time a migratory support role for this lineage in the mouse embryo. CH lineage tracing using *Wnt3a^iresCre^* further supports the astrocytic rather than multipotent radial glia nature of the scaffold because following an early neurogenic phase, when CR cells are produced, CH cells then give rise to the GFAP+ scaffold, but not to neurons or OPCs. In the spinal cord, *Aldh1l1*-GFP is first detected at E12.5^50^ and we observe a similar onset of expression of ALDH1L1 in the CH astrocytic progenitors, which we suggest may represent the fimbrial glioepithelium^22^. This is in contrast with what is observed elsewhere in the developing forebrain where astrocytes are known to arise around E16.5^51^, and suggests an earlier emergence in this region. Interestingly, CH derived CR cells are the first neurons generated in the brain^52^. Since the gliogenic switch is controlled by differentiating neurons^53^, it is tempting to speculate that the initial emergence of CR cells may explain early gliogenic induction in the CH. Finally, in the supragranular scaffold, ALDH1L1 and GFAP are both present but they do not colocalise. This supports a mixed origin for this part of the scaffold, and furthermore suggests a different cellular composition. In fact, exclusive expression of GFAP in some cells suggests that, in this domain, DNE-derived RGCs support neuronal migration, in addition to some CH-derived astrocytic progenitors.

Other migration cues are necessary for granule neuron guidance. CH-derived CR cells play an important role through the release of chemokines, such as Reelin^26^ and SDF1 (CXCL12)^54^. We did not observe any significant alteration in *Reelin* and *Cxcl12* expression indicating that CR cells are not affected by *Sox9* deletion. Therefore, our work highlights a crucial role for the fimbrial glial scaffold for 1ry-to-3ry matrix progenitor migration from E18.5, while the supragranular scaffold facilitate 3ry matrix cell distribution. Detailed investigations are needed to characterize how the fimbrial glial scaffold support progenitor migration.

### Role of SOX9 during DG morphogenesis

Impairment of DNE progenitor migration in *Sox9^fl/fl^;Wnt3a^iresCre/+^* mutants demonstrates that loss of SOX9 in these cells does not explain the migration defect. However, this does not rule out a role for SOX9 in DNE progenitors. Indeed, while we did not observe a significant alteration in progenitor emergence, differentiation and survival embryonically, there is a significant decrease in intermediate progenitor numbers early post-natally. This reduction coincides with formation of the ectopic cluster, where 35% of TBR2+ progenitors are unable to reach the developing DG in *Sox9^fl/fl^;Sox1^Cre/+^* mutants (Fig. 3.Ei). Their survival and expansion might be affected away from their destination, and account for the reduction in progenitor total numbers. While we did not detect a significant increase in apoptosis in the ectopic cluster at P2, these cells are absent in the adult brain, suggesting a post-natal loss. Alternatively, reduction of TBR2 cells could indicate that SOX9 is required for maintenance and/or expansion of migrating NSCs^14^, leading to post-natal reduction of a pool of newly formed TBR2 cells that was not detected by our analysis. In support of a role of for SOX9 in DNE cells, adult DG functionality is affected by embryonic deletion of SOX9, because we observe compromised memory formation abilities in *Sox9^fl/fl^;Sox1^Cre/+^* adults. This is the most straightforward explanation; however, we cannot exclude that other brain regions potentially affected by *Sox9* deletion may also impact on this impaired behaviour (e.g. altered locomotion, smell or sight). We^30^ and others^31, 55^ have indeed previously shown that SOX9 regulates progenitor formation and expansion. However, in other contexts, such a role was not observed^28, 32, 56, 57^. This variability may depend on the cellular context, but timing of the deletion is also relevant. Compensation by other members of the SOXE family, in particular SOX8, has been shown to explain recovery of some defects due to *Sox9* loss^58^. In our context, it is likely that both compensatory mechanisms and timing contribute to the difference in the severity of the defects observed after *Sox9* loss in different models. Analysis of SOX9/SOX8 double mutants would clarify this possibility. Finally, SOX9 is also expressed in adult DG NSCs^59^, where it is likely required for maintenance of SVZ NSCs^30^. As discussed above, compromised memory forming abilities are observed in *Sox9^fl/fl^;Sox1^Cre/+^* adults, which could well reflect the reduced numbers of neuronal progenitors reaching the DG. However, impaired adult neurogenesis could contribute to this phenotype, and although it is beyond the scope of the current study, this warrants further investigation. Moreover, as discussed above, SOX9 is likely to be required in DNE progenitors for formation of the supragranular scaffold, most likely as an inducer of gliogenic or RGCs fate^30, 32^.

The importance of SOX9 for the acquisition of gliogenic potential is demonstrated by the loss of the fimbrial scaffold in *Sox9^fl/fl^;Wnt3a^iresCre/+^* mutants. In the astrocytic lineage, SOX9 expression is maintained at high levels^60^ and it is required for astrocytic specification in the spinal cord^28^ and anterior CNS^30, 48, 55^. In the spinal cord it has been shown to induce expression of NFIA, with which it then interacts to activate expression of astrocytic genes^28^. In contrast, we show here that NF1A/B expression is not affected in the developing forebrain following *Sox9* deletion. Therefore other factors and pathways must be involved for induction of *Nf1a* expression, which could include BRN2^61^, WNT^62^ and/or NOTCH^63^. Furthermore, as shown in the spinal cord^28^, gliogenesis in CH is simply delayed in *Sox9^fl/fl^;Sox1^Cre/+^* and *Sox9^fl/fl^;Nestin-Cre* mutants. This could reflect a reduced transcriptional activity of NF1 factors without SOX9, as previously shown for NF1A^28^. As discussed above, recovery could also be due to compensatory expression of SOX8. Further analyses are required to understand the mechanisms underlying recovery of the fimbrial scaffold.

### Conclusions

The ability of the fimbrial glial scaffold to support progenitor migration is an exciting new finding. It will be important to characterize the molecular mechanism utilized by the scaffold for this function, in fact whether this support is based on release of chemoattractants and/or relies on cell-cell or cell-matrix interactions, is still unknown. We show that astrocytes forming the fimbrial scaffold are closely intermingled with migrating neuronal progenitors of the 2ry matrix, in fact they could form tubules around them similarly to how this same cell type supports neuroblast migration along the rostral migratory stream of the adult brain^7, 8^. This aspect of scaffold functionality could be addressed using *Aldh1l1-Cre*^50^ to delete ligands potentially involved.

In addition, our CH lineage tracing analysis using *Wnt3a^iresCre^;R26^ReYFP^* shows that progenitors in the CH exclusively generate CR cells and then switch to formation of the glial scaffold. However, we found around 6.5% of granule neurons in the Wnt3a lineage, suggesting that a proportion of neuronal progenitors may originate from the CH. This exciting possibility requires further confirmation with additional lineage tracing analyses; it also suggests that this subpopulation may have different characteristics. Whether other cells forming the adult DG, such as adult NSCs, also originate from the CH, is unknown and requires further investigation.

In conclusion, SOX9 plays sequential roles during CNS development as cells progress from a NSC fate, in which the protein is required for induction and also maintenance, to acquisition of gliogenic potential. Experimental manipulation of its expression levels highlights aspects of its function, according to the cellular context and also timing, which is presumably explained by the pattern of expression of redundant SOXE members, and its different interactors. Here we reveal that DG development is particularly sensitive to early loss of *Sox9* due to the failure to generate an astrocytic scaffold that aids neuronal migration. Extensive cell migration^64^ and progenitor pool expansion^65^ might underlay the vulnerability of this region, illustrated by the lasting consequences of transiently impaired cell migration.

## ACKNOWLEDGEMENTS

We are grateful for help and support from all past and present members of the Lovell-Badge’s lab. We thank François Guillemot (the Francis Crick Institute) for helpful discussions. Distance.gui software for analysis of granule neurons distribution within the forming DG was written by Dr. Olivier Pierre Friard and kindly donated by Prof. Federico Luzzati (Neuroscience Institute Cavalieri Ottolenghi – NICO). We also thank Dr. Lucas Baltussen for providing training for *in utero* electroporation and kind donation of pCAG-hyPBase and pPB-CAG-DsRed plasmids. We thank Dr Celia Garau and Dr Richard Lilley for assistance in setting up the behavioural test and analysis. Finally, we also thank Biological Services, Experimental Histopathology, Advanced light microscopy and Mechanical Engineering platforms at the Francis Crick Institute, for their excellent assistance and technical support and the Mutant Mouse Regional Resource Center U42OD010918 for providing aliquots of *Wnt3a^iresCre^* cryo-preserved sperm.

This work was supported by the Medical Research Council, U.K. (U117512772, U117562207 and U117570590) and the Francis Crick Institute which receives its core funding from Cancer Research UK (FC001107), the UK Medical Research Council (FC001107), and the Wellcome Trust (FC001107).

## MATERIAL AND METHODS

### Mouse strains, husbandry and genotyping

All experiments carried out on mice were approved under the UK Animal (scientific procedures) Act 1986 (Project license n. 80/2405 and PP8826065). Mouse husbandry, breeding, ear biopsies and vaginal plug (VP) checks were performed by the Biological Research Facility team of the Francis Crick Institute. Animals were kept in individually ventilated cages (ICV) with access to food and water *ad libitum*. The VP day was considered as 0.5 day from time of conception (E0.5) and the day of birth termed P0. All mouse lines used were previously described: *Sox9^fl/fl^* conditional targeted mutation, MGI: 2429649^36^; *Sox1^Cre/+^* targeted mutation, MGI: 3807952^34^; *Nestin-Cre* transgenic mutation, MGI: 2176173^33^; *Wnt3a^IRES-Cre^* targeted mutation, MGI: 98956^17^; *R26R^eYFP^* targeted mutation, MGI: 2449038^66^. To obtain *Sox9* conditional mutations, *Sox9^fl/fl^* mice were crossed with either *Sox1^Cre/+^*, *Nestin-Cre* or *Wnt3a^IRES-Cre^* mice. All Cre lines were kept in heterozygosity. To verify Cre recombination pattern, *R26R^eYFP^* reporter allele was also present in all samples analysed, even when not indicated. Genotyping of embryos and adult mice was performed by Transnetyx.

### Behavioural analysis

The novel object recognition test (NORT) was used to analyse memory formation in adult mice, as the ability to discern between new and familiar objects. A 40×40cm arena made of white Plexiglas was built by the Francis Crick Institute mechanical engineering facility. Pairs of different objects were switch between cohorts of animals to avoid biases due to object conformation. Logitech 910C webcam and software were used to record behavioural tests and videos were used for analyses. Mice were acclimatised to the testing room for 1 hour before starting the test. The operator was alone with the mice during the duration of the test to avoid any disturbance. The behavioural test was performed on three consecutive days, and each mouse spends 5min in the arena per day. On the first day, mice were placed in an empty arena for adaptation. On the second day, mice were exposed to two identical objects, for training. On the third final day, mice were confronted to one familiar object (used the previous day) and a new, different object. Arena and objects were disinfected and rinsed before every run. Objects location within the arena was consistent among animals. The recordings from the first day of NORT (adaptation day) were used to perform open field test. Ethovision XT (Noldus) was used for the analysis of video recordings. For NORT, in each video, a circular area of 4cm radius around each object was considered as object “exploration area”. Time spent in this area was considered as “exploration time”. Mice that did not explore both objects on the second day were excluded from further analysis. For the open field test, the arena centre was considered as a central square area 4cm apart from the arena borders. Time spent in this area was considered as “time spend it centre” and was calculated with Ethovision XT.

### EdU injection

For cell birth-dating experiments, 10mg/ml EdU solution, from Click-iT^TM^ EdU imaging kit, was injected intraperitoneally in pregnant females at a dose of 30µg/g body weight. Injection was performed at either E16.5 or E18.5 stage of pregnancy and samples collected at E18.5 or P2 respectively. Schematic of protocols in Fig. S.6.E,I.

### Tissue harvesting and staining

Pregnant females were killed by cervical dislocation, embryonic heads were dissected out in chilled PBS and fixed by immersion in chilled 4% PFA at 4°C for 1 to 2 hours. From E16.5 onwards, brains were dissected out of the skull before fixation. P2 pups were killed by cervical dislocation, brains were dissected out on chilled PBS and fixed by immersion in chilled 4% PFA at 4°C for 2 hours. After fixation, embryonic brains were washed once in PBS, cryopreserved in sucrose 30%, then embedded in OCT, frozen on dry ice and stored at −80°C. Samples were cryosectioned at 14μm and sections placed on Superfrost Plus glass slides, air dried for 5 minutes, washed twice in PBS for 5 minutes. For some antibodies (listed in Table 1), antigen retrieval was performed immerging slides in 10% target retrieval solution pH6.1 diluted 1:10 in distilled water, for 30 minutes at 65°C or 15 minutes at 95°C, then washing twice in PBS for 10 minutes. Slides were then incubated in a humidified chamber in blocking solution (10% donkey serum in 0.1% triton X-100 PBS) for at least 30 minutes, then incubated overnight at 4°C with primary antibodies diluted in blocking solution (dilution indicated in Table 1). The following day, sections were washed twice for 5 minutes in 0.1% triton X-100 PBS, incubated with secondary antibodies (Table 2) and DAPI diluted in blocking solution for 2 hours at room temperature in a dark humified chamber. Finally, sections were washed again twice for 5 minutes in PBS, briefly in distilled water, air dried and coverslip mounted with Aqua-poly/Mount. Apoptosis detection with terminal deoxynucleotidyl transferase dUTP nick end labelling (TUNEL) assay was performed using the ApopTag® Red *In Situ* Apoptosis Detection Kit, following manufacturer’s instructions, after secondary antibody incubation. Detection of DNA-incorporated EdU was performed using Click-iT^TM^ EdU imaging kit, following manufacturer’s instructions, after secondary antibody incubation.

**Table 1:**
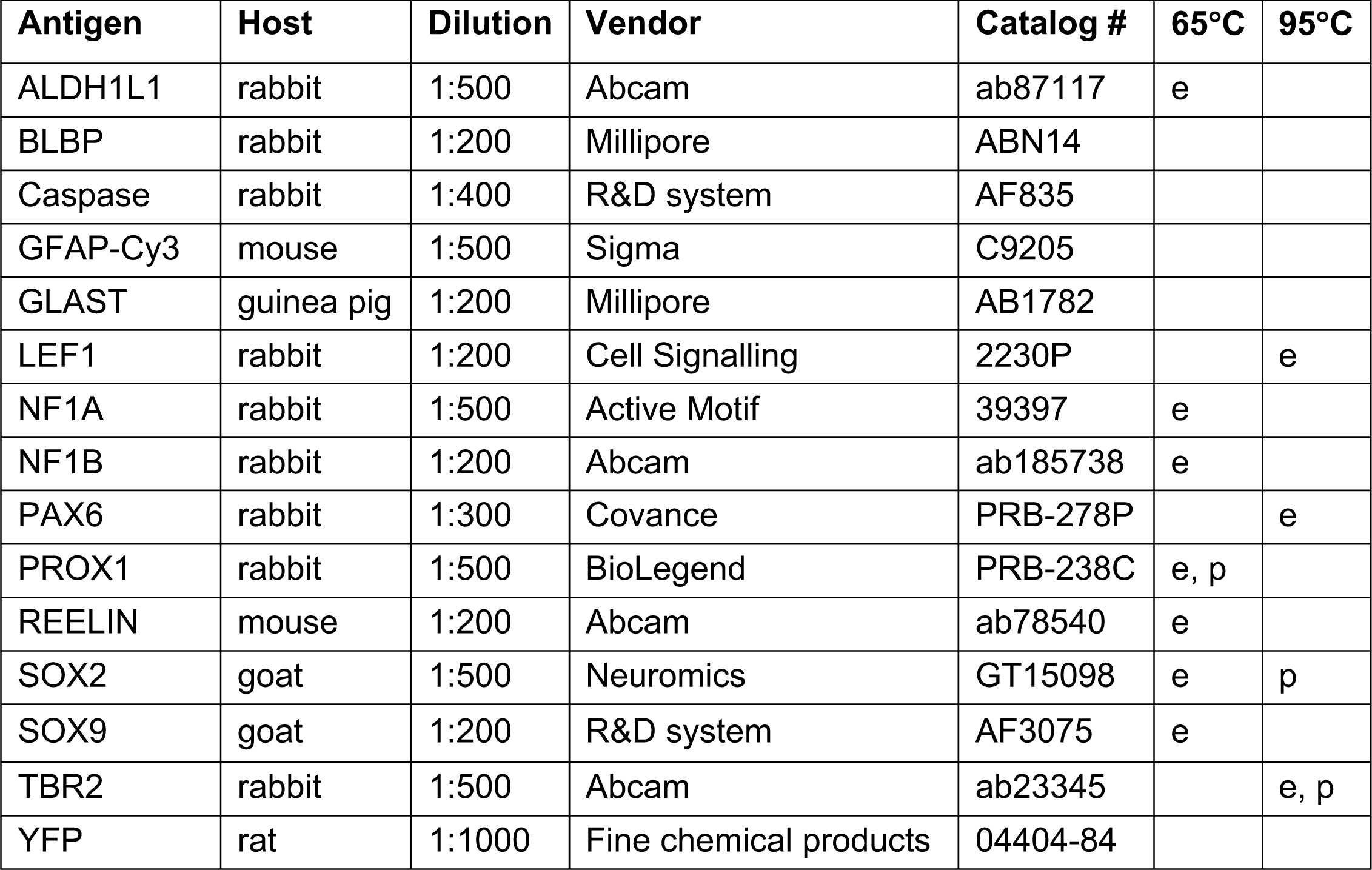
List of primary antibodies used. Antigen retrieval protocol (30 minutes in 65°C waterbath or 15 minutes in 95°C decloaking chamber) was performed for the indicated samples (e: embryos; p: pups).

| Antigen | Host | Dilution | Vendor | Catalog # | 65°C | 95°C |
| --- | --- | --- | --- | --- | --- | --- |
| ALDH1L1 | rabbit | 1:500 | Abcam | ab87117 | e |  |
| BLBP | rabbit | 1:200 | Millipore | ABN14 |  |  |
| Caspase | rabbit | 1:400 | R&D system | AF835 |  |  |
| GFAP-Cy3 | mouse | 1:500 | Sigma | C9205 |  |  |
| GLAST | guinea pig | 1:200 | Millipore | AB1782 |  |  |
| LEF1 | rabbit | 1:200 | Cell Signalling | 2230P |  | e |
| NF1A | rabbit | 1:500 | Active Motif | 39397 | e |  |
| NF1B | rabbit | 1:200 | Abcam | ab185738 | e |  |
| PAX6 | rabbit | 1:300 | Covance | PRB-278P |  | e |
| PROX1 | rabbit | 1:500 | BioLegend | PRB-238C | e, p |  |
| REELIN | mouse | 1:200 | Abcam | ab78540 | e |  |
| SOX2 | goat | 1:500 | Neuromics | GT15098 | e | p |
| SOX9 | goat | 1:200 | R&D system | AF3075 | e |  |
| TBR2 | rabbit | 1:500 | Abcam | ab23345 |  | e, p |
| YFP | rat | 1:1000 | Fine chemical products | 04404-84 |  |  |

**Table 2:**
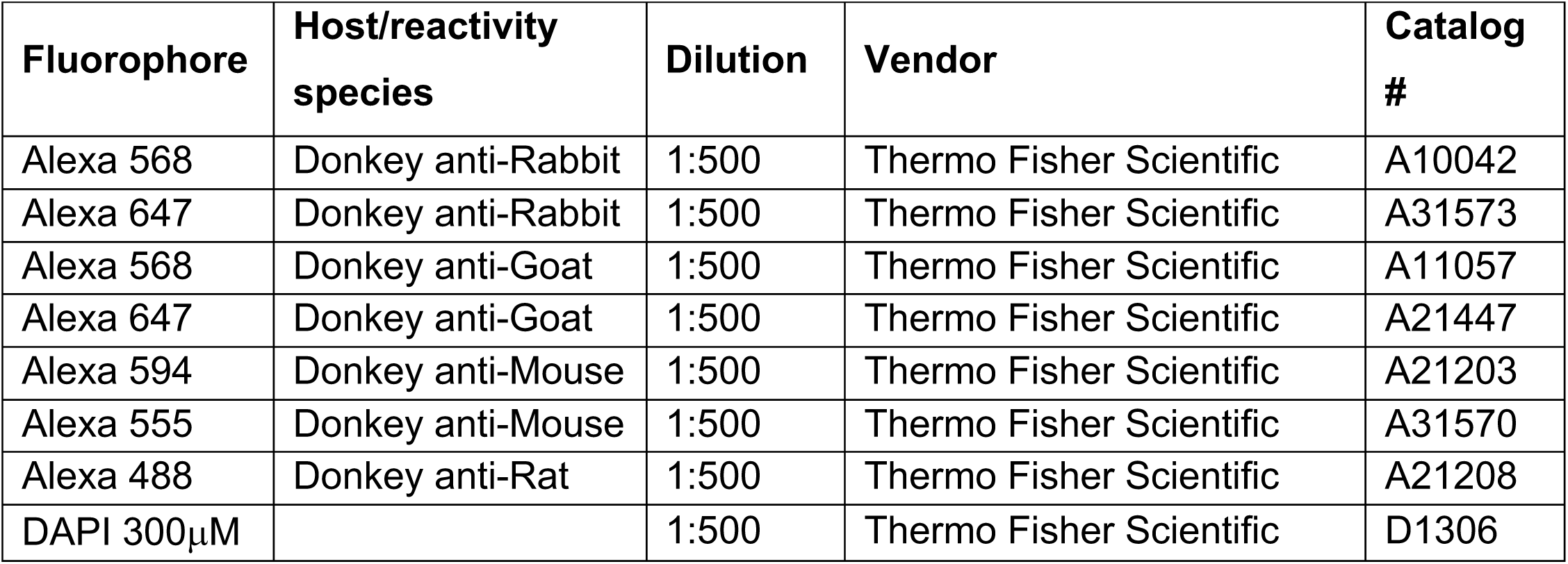
List of secondary antibodies and nuclear staining used.

| Fluorophore | Host/reactivity species | Dilution | Vendor | Catalog # |
| --- | --- | --- | --- | --- |
| Alexa 568 | Donkey anti-Rabbit | 1:500 | Thermo Fisher Scientific | A10042 |
| Alexa 647 | Donkey anti-Rabbit | 1:500 | Thermo Fisher Scientific | A31573 |
| Alexa 568 | Donkey anti-Goat | 1:500 | Thermo Fisher Scientific | A11057 |
| Alexa 647 | Donkey anti-Goat | 1:500 | Thermo Fisher Scientific | A21447 |
| Alexa 594 | Donkey anti-Mouse | 1:500 | Thermo Fisher Scientific | A21203 |
| Alexa 555 | Donkey anti-Mouse | 1:500 | Thermo Fisher Scientific | A31570 |
| Alexa 488 | Donkey anti-Rat | 1:500 | Thermo Fisher Scientific | A21208 |
| DAPI 300μM | | 1:500 | Thermo Fisher Scientific | D1306 |

For 3D reconstruction, samples were cryosectioned at 50μm and sections placed floating in a 24 well plate in PBS. Sections were washed, processed for antigen retrieval at 65C and incubated in blocking solution (10% donkey serum in 0.5% triton X-100 PBS) as described above. Primary and secondary antibody incubation (antibodies diluted in 10% donkey serum in 0.5% triton X-100 PBS; dilution indicated in Table 1 and 2) was respectively 3 and 1 nights. Floating sections were then mounted on Superfrost Plus glass slides, air dried and coverslip mounted with Aqua-poly/Mount.

For hematoxylin and eosin (H&E) staining, pregnant females and adult mice were killed by cervical dislocation, embryonic and adult brains were dissected out in chilled PBS, fixed overnight in Bouin’s solution at 4°C, washed twice for 10 minutes in 70% ethanol and stored in 70% ethanol until processing. Samples were embedded in wax, sectioned at 4µm and stained by the Francis Crick Institute Experimental Histopathology facility.

### *In situ* hybridization probe formation and staining

Digoxigenin (DIG)-tagged antisense RNA probes were made from an ampicillin-resistant plasmid (kindly gifted by Dr. Paul Sharp) containing *Sox9* cDNA followed by a T7 promoter. Plasmid was amplified with *E. Coli* culture and purified with NucleoBond Xtra Midi plus kit. 5µl of plasmid (corresponding to 5-10µg) was linearized with 1µl of SmaI enzyme, 2ul of 10x SmartCut buffer and 12ul of RNase free water and confirmed with 1% agarose gel electrophoresis. Linearized plasmids were purified by phenol-chloroform extraction and precipitated by adding 1µl glycogen, 1/20 of sodium acetate 3M and equal volume of 100% ethanol, incubated at −20°C for 1 hour. The precipitate was recovered by centrifugation at 13000 RPM at 4°C for 15min, air dried and resuspended in 16µl of RNase free water. For DIG-tagged probes synthesis, 1µl linearized plasmid, 1X transcription buffer, 2µl of DIG-tagged nucleotides, 1µl T7 RNA polymerase, 0.5µl RNase inhibitor, 1µl DTT 100mM and 11.5µl of RNase free water were incubated at 37°C for 2 hours. Probe formation was confirmed with 1% agarose gel electrophoresis. Probes were precipitated by adding 1µl glycogen, 8µl lithium chloride 5M, 2/3 of final volume of 100% ethanol and 1/3 of final volume of RNase free water, incubated at −80°C for 30 minutes. Precipitates were recovered by centrifugation at 13000 RPM at 4°C for 15 minutes, then washed in 70% ethanol (v/v) centrifuged at 13000 RPM for 15min at 4°C. Pellet containing the RNA probes were then air-dried at 37°C and resuspended in 50-100µl of ISH hybridization buffer (50% (v/v) deionised formamide, 4X SSC, 0.01M *β*-mercaptoethanol, 10% dextran sulphate, 2X Denhart’s solution, 0.23mg/ml yeast t-RNA diluted in RNase free water).

For *in situ* hybridisation (*ISH*) staining, cryosections were initially air-dried, washed twice for 5 minutes in PBS, fixed 30 minutes with 4% PFA at room temperature in a humidified chamber, washed twice for 10 minutes in PBS, and incubated in pre-hybridization buffer (50% (v/v) deionised formamide, 1X saline-sodium citrate (SSC) diluted in RNase free water) for 1 hour in a 65°C waterbath. For each slide, 1μl of RNA probe was denaturated in 200μl hybridization buffer for 10 minutes at 70°C, then applied on sections. Hybridization was carried out in a humidified chamber overnight in a 65°C water bath. The following day, sections were washed twice for 15 minutes, and once for 30 minutes with pre-warmed washing buffer (50% (v/v) deionised formamide, 1X SSC, 0.1% (v/v) Tween 20 diluted in MilliQ water) at 65°C, then twice for 30 minutes with MABT (20mM Maleic acid, 30mM sodium chloride (NaCl), 0.02% (v/v) Tween 20, adjusted to pH7.5 and diluted in MilliQ water) at room temperature. Sections were then incubated in a humidified chamber at room temperature for 1 hour in blocking buffer (10% (v/v) sheep serum, 2% (v/v) blocking reagent diluted in MABT), then overnight in anti-DIG coupled to alkaline phosphatase antibody (*α*-DIG-AP) diluted 1:1500 in blocking buffer. The following day, sections were washed four times for 20 minutes at room temperature with MABT then twice for 10 minutes in pre-staining solution (100mM NaCl, 50mM magnesium chloride (MgCl2), 100mM Tris-HCl pH9.5, 1% (v/v) Tween 20 in MilliQ water). Alkaline phosphatase staining was then performed incubating slides with staining solution (pre-staining solution plus 5% (w/v) Polyvinyl alcohol (PVA), 4.5µl of Nitrotetrazolium Blue chloride (NBT) and 3.5µl of 5-bromo-4-chloro-3-indolyl-phosphate (BCIP) per ml of staining solution) for 1 hours to 2 days at 37°C in a dark chamber. Once the desired staining intensity was reached, the reaction was stopped washing sections twice for 10 minutes in 0.1% triton X-100 PBS. Finally, sections were fixed for 10 minutes in 4% PFA, washed twice for 5 minutes in PBS then briefly in distilled water, air-dried, and coverslip mounted with Aqua-poly/Mount.

### Quantitative PCR and analysis

For gene expression analysis, E12.5 embryos were dissected in sterile PBS, dorsal telencephalon and archicortex were separately snap-frozen in liquid nitrogen. RNA was extracted with the RNeasy Plus Micro kit, following manufacturer’s instructions. Complimentary (c) DNA was synthetized from 250ng of extracted RNA in 1X qScript cDNA SuperMix diluted in RNase free water and incubated on the Tetrad 2 Thermal Cycler following the indicated reverse transcription protocol. Resulting cDNA was diluted in RNase free water at a final concentration of 200µg/µl. Transcripts were quantified with quantitative PCR (qPCR), mixing 800µg of cDNA in 1X ABsolute QPCR SYBR Green ROX Mix and 40nM of primer mix (forward and reverse primers were pre-mixed; primers sequences are indicated in Table 3). Expression level of the house-keeping gene Glyceraldehyde 3-phosphate dehydrogenase (GAPDH) was used as reference. Each sample was run in technical triplicates, in case of high variability, one of the technical triplicates was removed. Number of biological replicates are indicated for each experiment (n).

**Table 3:**
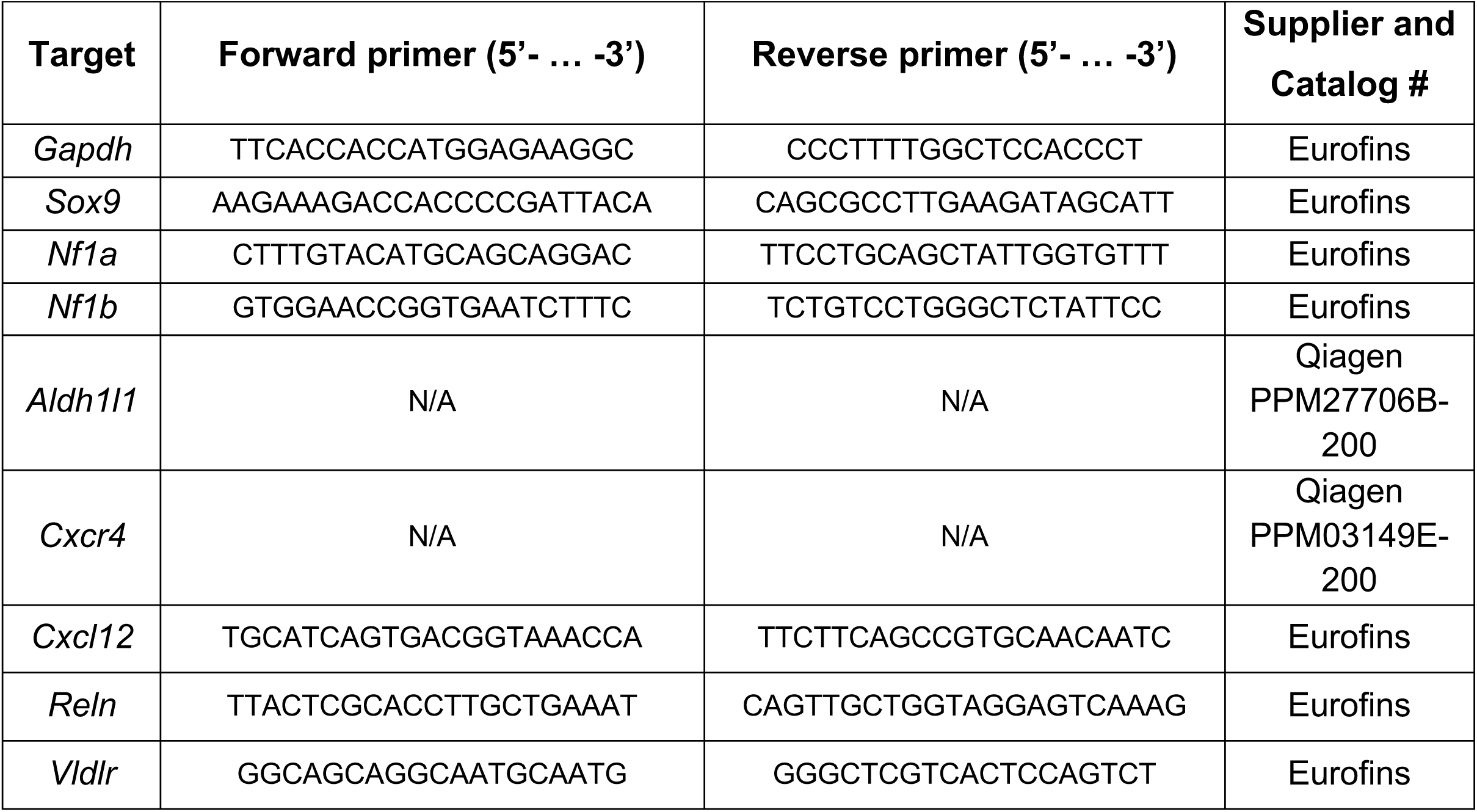
List of primers used for qPCR.

Relative expression of the genes of interest were calculated by normalisation of the detected expression value to the geometric mean of the reference GAPDH gene using the *ΔΔ*Ct method^67^. More precisely, average cycle threshold (Avg Ct) was first calculated among technical triplicates or duplicates of each sample. Average delta Ct (Avg *Δ*Ct) was then deduced by subtracting GAPDH Avg Ct to sample Avg Ct. The relative quantification (RQ) of cDNA for each gene was calculated as 2^-AvgΔCt^. The fold change of each sample was calculated in reference to the average RQ of control samples group (control RQ) as: sample RQ/control RQ. The qPCR final results are shown as histograms, where each bar shows the average fold change of experimental replicates. Error bars are represented as standard error of the mean (SEM).

### *In utero* electroporation

For *in utero* electroporation (IUE), the *piggyBac* transposon system was used to avoid episomal plasmid loss upon cell division. pCAG-hyPBase and pPB-CAG-DsRed plasmids were kindly donated by Dr. Lucas Baltussen and have been previously described^68^. Plasmids was amplified with *E. Coli* culture, purified with EndoFree Plasmid Maxi kit and mixed together at a concentration of 1µg/µl per plasmid, with 0.05% of FastGreen in injectable water.

IUE was performed on E15.5 embryos. One before surgery, analgesia (Carprofen, dose 50mg/ml) was administered via drinking water to single caged pregnant females. On surgery day, pregnant females were anaesthetised using isoflurane and subcutaneous injection (10mg/kg of meloxicam and 0.1mg/kg of buprenorphine in injectable water). Females’ eyes were kept moist using Viscotears eye gel. Anesthetised female was shaved on the abdomen, cleaned with chlorhexidine and moved to surgical area, where body temperature was monitored on a heating pad. Laparotomy and exteriorisation of the uterus were then performed. 10µl of DNA was loaded using micro loader tips into Ethylene Oxide (EtO) gas sterilised borosilicate glass capillaries (1.0 OD x 0.58 ID x 100L mm) which were pulled with a micropipette puller and the tip was broken using forceps. 1µl of solution containing the plasmid DNA was injected into the lateral ventricle of each embryos using a Femtojet pico dispenser, followed by electroporation (5 pulses 38V of 50ms with 1s interval) with EtO gas sterilised 5mm paddle type electrodes. The uterus was gently reinserted into the abdomen, then abdominal wall and skin were sutured separately. Mice were placed in a recovery chamber for a few hours. Analgesia (Carprofen, dose 50mg/ml) was administrated in drinking water for the following 48 hours. Electroporated embryos were harvested at P2.

### Software for cell migration analysis

Distance.gui software was used to analyse PROX1+ cells distribution within the forming DG. It was written by Dr. Vivien Labat-gest and kindly donated by Prof. Federico Luzzati. The software calculates the distance in pixels between a point and line, which in this case are single PROX1+ cells and their migration line, respectively (schematic in Fig. 2.F). Therefore, the software input files for each image are the ImageJ cell counter result *.xml* file (representing PROX1+ cells points coordinates) and the XY coordinates of the reference line extracted from ImageJ as a *.txt* file (representing the migration line). The output file is a *.txt* file containing a list of numbers representing the distance in pixel of each PROX1+ cell from the migration line. For each picture, the range of PROX1+ cell distribution (most distant cell from the migration line) was used to divide the forming DG area in 10 bins, then percentage of PROX1+ cell per bin was calculated and plotted as a line (Fig. 2.F and 5.I). Cells in bin #1-5 would be closer to the migration line, therefore representing the lower DG blade, compared to cells in bin #6-10, representing the upper DG blade.

### Statistical and image analysis

H&E and *ISH* stained sections were acquired with Leica DM750 light microscope, and LAS (Leica Application Suite) EZ software was used for acquisition. Sections processed for EdU, TUNEL and immunofluorescence staining were imaged using Leica TCS SPE confocal microscope with 10x, 20x and 40x objectives. LAS AF software was used for acquisition. Acquisitions were performed as 1.5µm Z-stacks, with bidirectional X. For image analysis and counting, ImageJ and QuPath softwares were used. For cell quantification, 5 different images were acquired and counted per analysed area for each sample. For 3D reconstructions, acquisitions were performed as 1µm Z-stacks, with bidirectional X and IMARIS was used image processing.

Statistical analysis of cell number quantification and qPCR analysis was performed on Prism 7 (Graphpad), calculating student’s two-sided unpaired *t-tests*, when comparing two groups, or ordinary one-way ANOVA, when comparing three or more groups. Analyses were performed parametrically, upon confirmation of samples normal distribution (on Prism). When the majority of sample groups within one analysis are not normally distributed (2 out of 3), statistical analysis was performed with Kruskal-Wallis test.

Histograms represent average quantification from the indicated number of biological replicates (n, minimum 3). Error bars for cell number quantification represent standard deviation (SD). Error bars for qPCR fold change analysis represent standard error of the mean (SEM). Data shown as percentage were processed with angular transformation before statistical analysis. P value is indicated as: ns:p>0.05; *:p≤0.05; **:p≤0.01; ***:p≤0.001; ****:p≤0.0001.

---

**Figure S.1.**
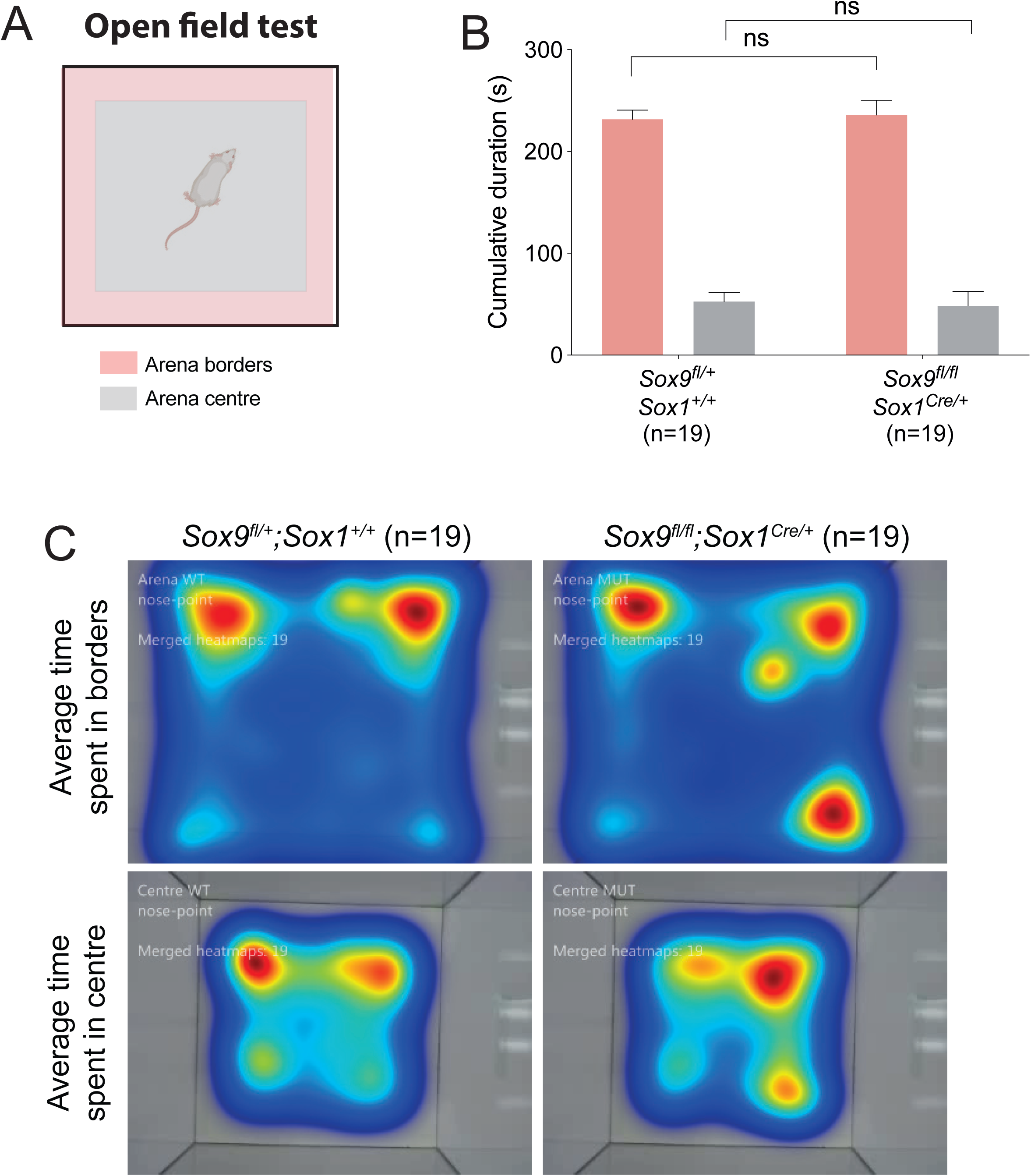
*Sox9^fl/fl^;Sox1^Cre/+^* adults do not show anxiety-like behaviour in open field test. (**A**) Schematic of arena subdivision between centre (grey) and borders (pink) used for open field test, performed to analyse anxiety-like behaviours. **(B,C)** Quantification of borders and centre of arena exploration time as cumulative seconds (**B**) and average time represented with heatmaps (**C**). We found no difference between *Sox9^fl/+^;Sox1^+/+^* and *Sox9^fl/fl^;Sox1^Cre/+^* mutant mice, both in time spent in the borders (231.72±8.78s and 235.93±14.38s, *t*-test P=0.2829) and in the centre (52.59±9.01s and 48.40±14.25s, *t*-test P=0.2864) of the arena. Therefore *Sox9^fl/fl^;Sox1^Cre/+^* mutant mice do not display anxiety-like behaviour in this test.

**Figure S.2.**
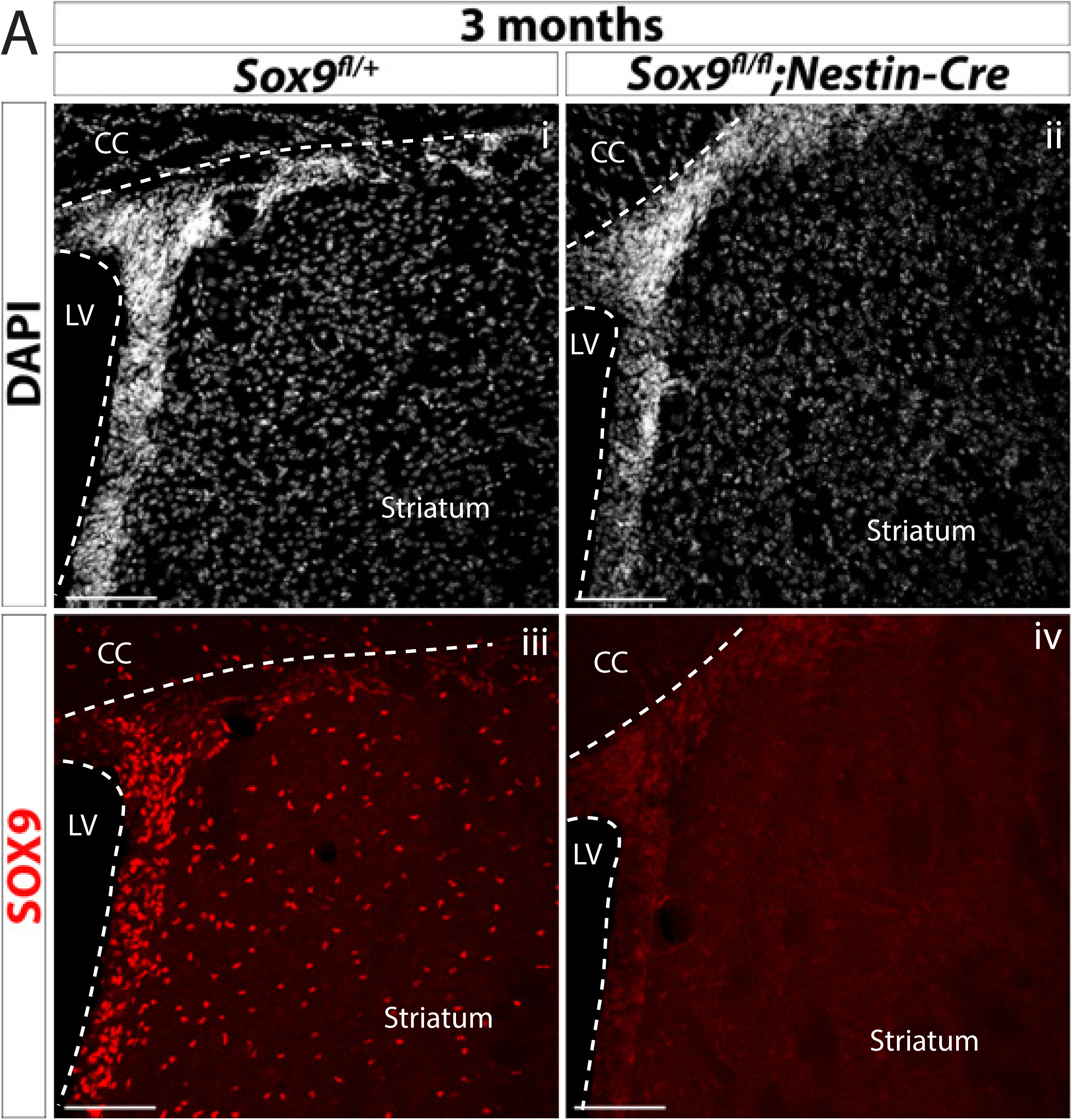
Absence of SOX9 expression in a 3 months-old *Sox9^fl/fl^;Nestin-Cre* mutant brain. (**A**) Immunofluorescence for SOX9 was performed to confirm absence of SOX9 expression in the brain of adult *Sox9^fl/fl^;Nestin-Cre* mutants (ii, iv) compared to controls (i, iii). LV: lateral ventricle; CC: corpus callosum. Scale bars represent 100µm.

**Figure S.3.**
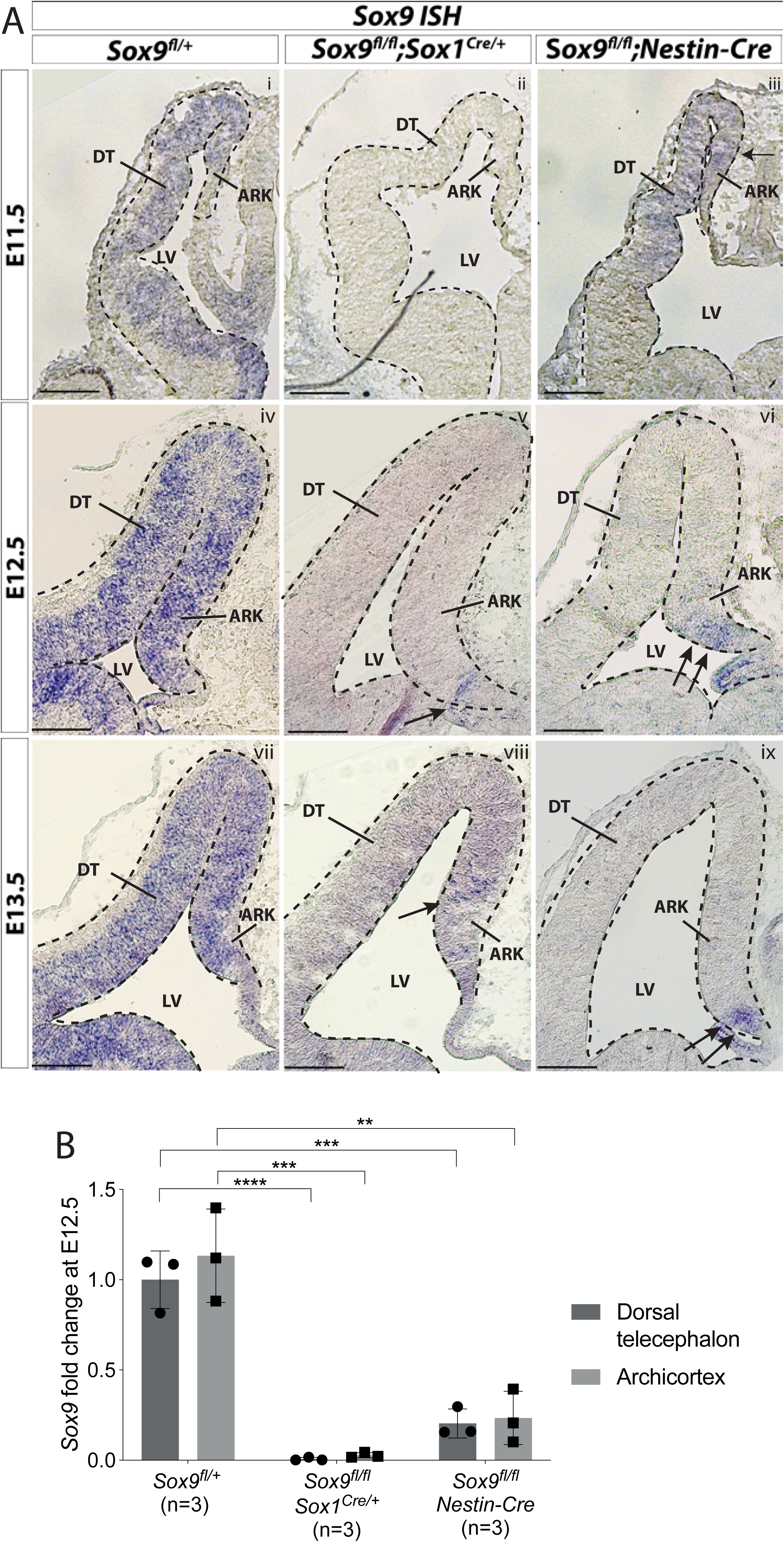
Qualitative and quantitative analyses of *Sox9* transcripts confirm residual *Sox9* expression in embryonic *Sox9^fl/fl^;Nestin-Cre* forebrains. **(A)** *ISH* for *Sox9* on *Sox9^fl/+^* (i,iv,vii)*, Sox9^fl/fl^;Nestin-Cre* (ii,v,viii) and *Sox9^fl/fl^;Sox1^Cre/+^*(iii,vi,ix) forebrains at E11.5 (i,ii,iii), E12.5 (iv,v,vi) and E13.5 (vii,viii,ix). Analysis of *Sox9* expression confirms ventral to dorsal activity of *Nestin-Cre* as *Sox9* is still expressed in the DT at E11.5 in *Sox9^fl/fl^;Nestin-Cre* mutants, (iii) and in the most ventral part of the ARK at E12.5 and E13.5 (arrows in vi, ix). Some rare cells still express *Sox9* in the archicortex of *Sox9^fl/fl^;Sox1^Cre/+^* mutants (arrow in v, viii). **(B)** Quantification of *Sox9* expression via qPCR from dissected DT and ARK of controls, *Sox9^fl/fl^;Sox1^Cre/+^* and *Sox9^fl/fl^;Nestin-Cre* E12.5 embryos. *Sox9* expression is drastically reduced in both DT and ARK of *Sox9^fl/fl^;Sox1^Cre/+^* compared to controls (*Sox9* fold change in DT: 0.006±0.008, P<0.0001; and ARK: 0.027±0.014, P<0.0001 Turkey’s multiple comparison test; ANOVA P<0.0001). *Sox9* expression was also significantly reduced in *Sox9^fl/fl^;Nestin-Cre* mutants compared to controls (*Sox9* fold change in DT: 0.204±0.08, P=0.0002; and ARK: 0.234±0.147, P=0.0017 Turkey’s multiple comparison test; ANOVA P=0.0005). *Sox9* tended to be present at higher levels in *Sox9^fl/fl^;Nestin-Cre* compared to *Sox9^fl/fl^;Sox1^Cre/+^*, which may reflect differences in Cre activities, however this was not statistically significant. LV: lateral ventricle; DT: dorsal telencephalon; ARK: archicortex. Scale bars represent 200µm.

**Figure S.4.**
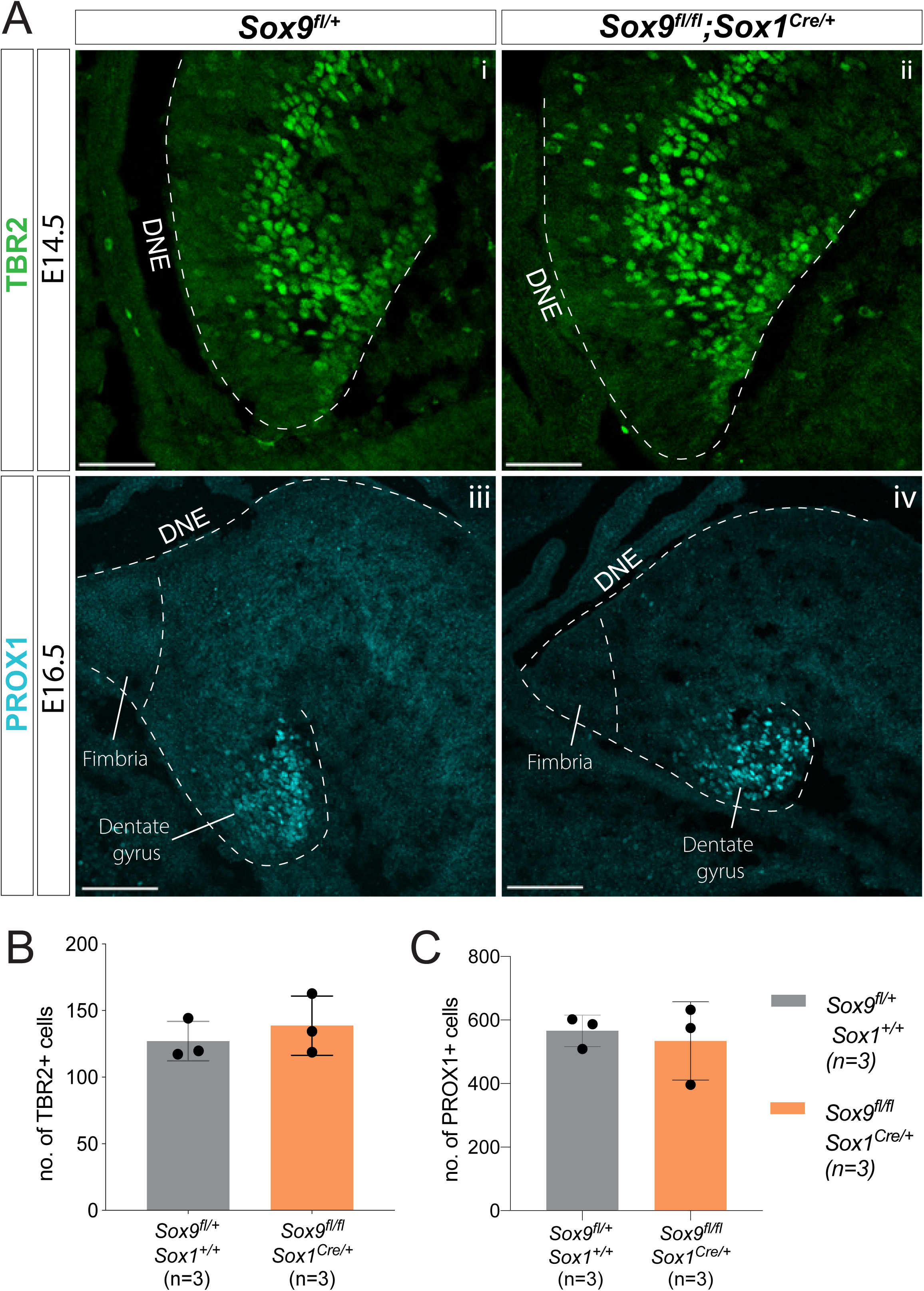
Initial emergence of intermediate progenitors and differentiating granule neurons is not affected by *Sox9* deletion. **(A)** Immunostaining for TBR2 (Ai-ii) and PROX1 (Aiii-iv) on respectively E14.5 and E16.5 controls and *Sox9^fl/fl^;Sox1^Cre/+^* mutant embryos. **(B-C)** Quantification shows that the total number of TBR2+ intermediate progenitors **(B)** and PROX1+ differentiating granule neurons **(C)** is similar in *Sox9^fl/fl^;Sox1^Cre/+^* mutants and controls. DNE: dentate neuroepithelium. Scale bars represent 50µm in (Ai-ii); 100µm in (Aiii-iv).

**Figure S.5.**
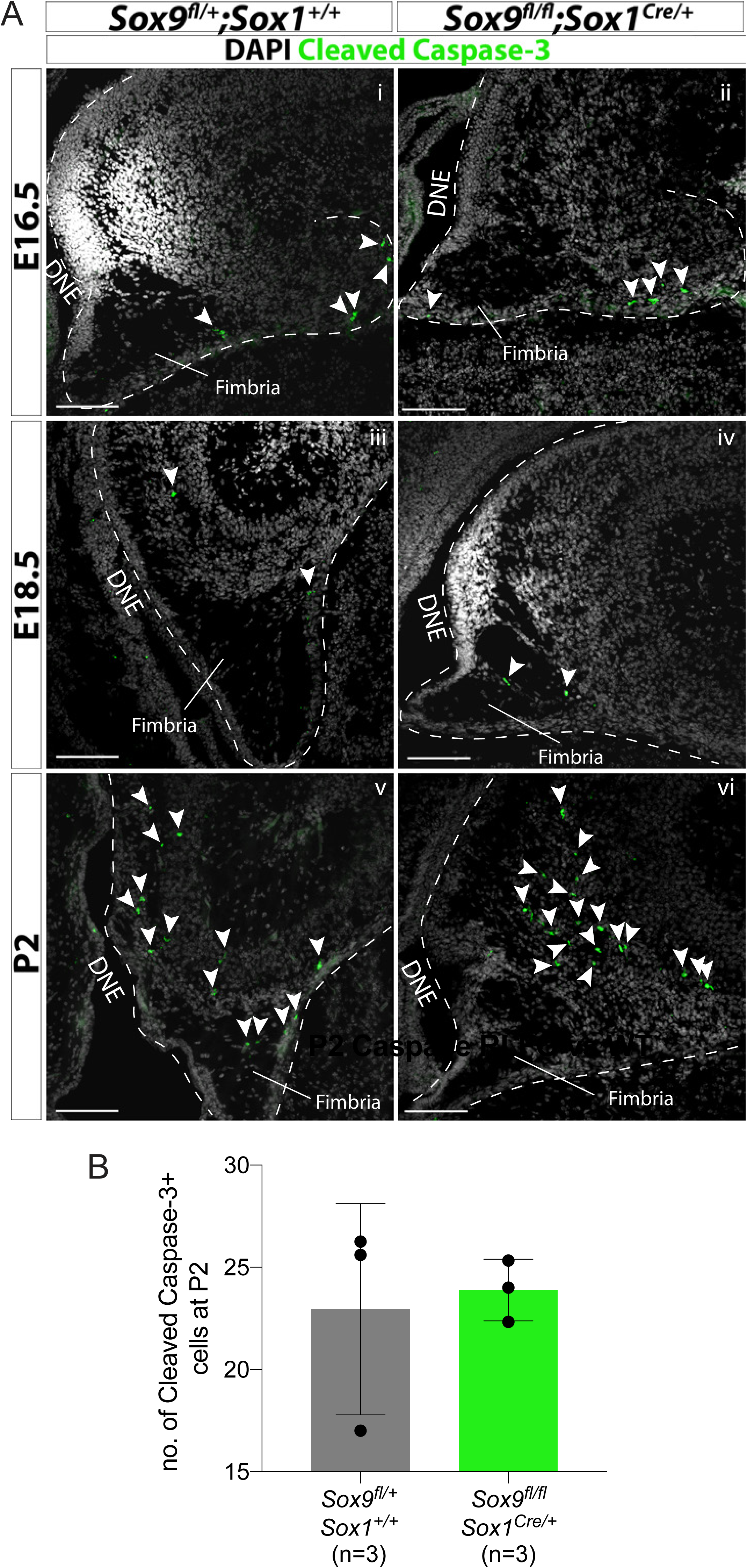
*Sox9* deletion is not associated with increased cell death in the developing DG. **(A)** Immunofluorescence for cleaved Caspase-3 at consecutive stages of DG development (E16.5: i, ii; E18.5: iii, iv; P2: v, vi) in *Sox9^fl/fl^;Sox1^Cre/+^* mutants compared to controls (arrowheads indicates cleaved Caspase-3+ cells). **(B)** Quantification of cleaved Caspase-3+ cells in 1ry and 2ry matrix of P2 pups shows that a similar number of apoptotic cells are present in *Sox9^fl/fl^;Sox1^Cre/+^* mutants compared to controls. DNE: dentate neuroepithelium. Scale bars represent 100µm.

**Figure S.6.**
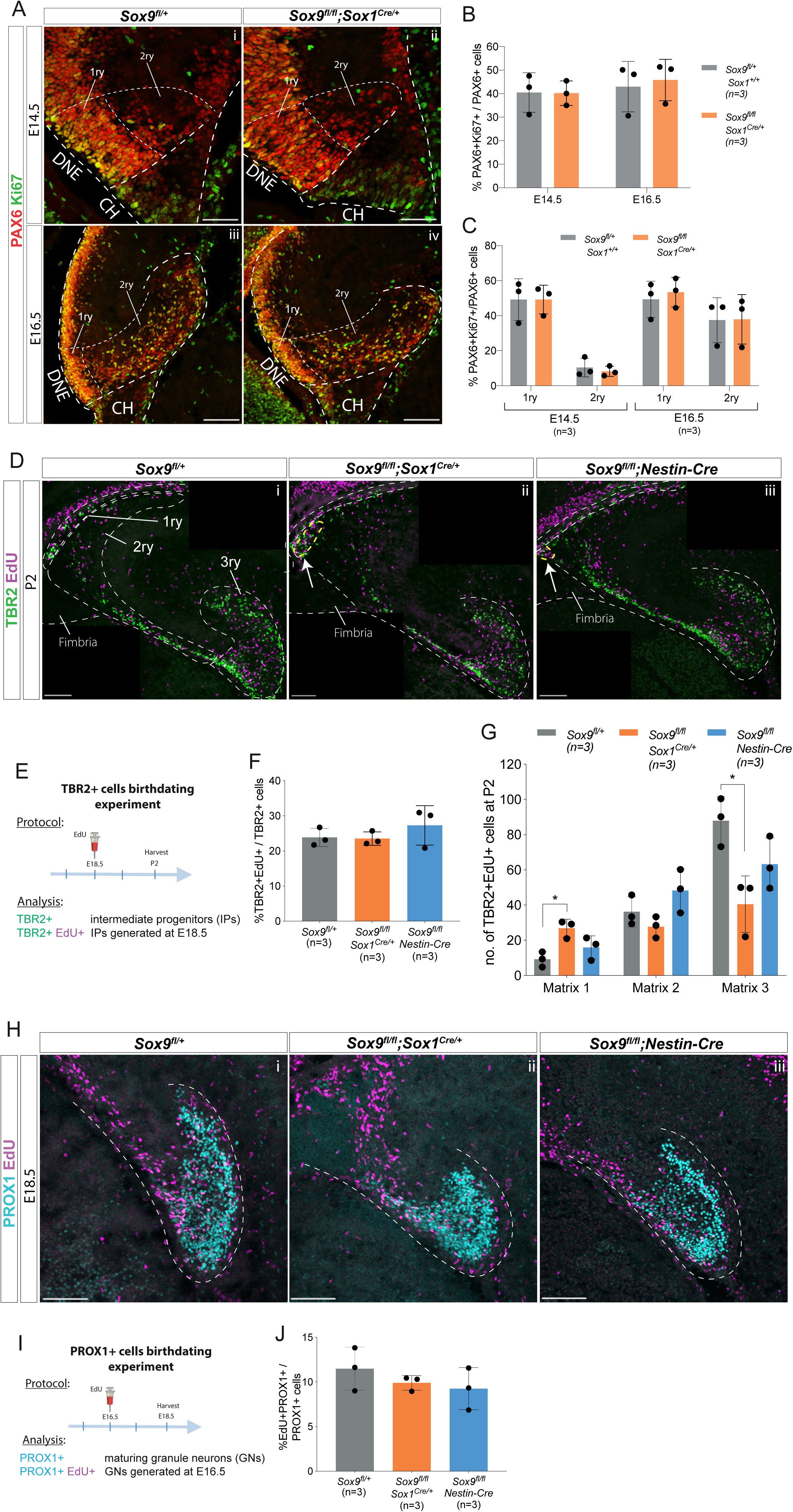
*Sox9* deletion does not alter rate of neural progenitor proliferation, emergence or differentiation towards a granule neuron fate. **(A-C)** Analysis of proliferation in early DG progenitors. (**A**) Double immunostaining for PAX6 and Ki67 on E14.5 (i-ii) and E16.5 (iii-iv) control and *Sox9^fl/fl^;Sox1^Cre/+^* mutant embryos. No difference was found in PAX6+ progenitor proliferation either in the total population, shown as % PAX6+Ki67+ on total PAX6+ cells (**B**), as well as analyzing 1ry and 2ry matrices separately (**C**). The regions considered for 1ry and 2ry matrix are indicated by the white dotted lines in (**A**). **(D-G)** DG progenitors EdU birth-dating experiment. (**D**) Double staining for TBR2 and EdU on P2 control, *Sox9^fl/fl^;Sox1^Cre/+^* and *Sox9^fl/fl^;Nestin-Cre* mutant brains. Arrows indicate accumulation of cells close to the ventricle in *Sox9* mutants. (**E**) Experimental protocol: EdU was injected at E18.5, samples harvested at P2 and immunostained for TBR2 and EdU. (**F**) TBR2+ progenitor emergence, quantified as the percentage of TBR2+EdU+/TBR2, was not affected by *Sox9* deletion. (**G**) Distribution of newly generated TBR2+EdU+ progenitor along the three matrices (as delineated in **D**i). A significant accumulation of TBR2+EdU+ newly formed progenitors was observed in the 1ry matrix of *Sox9^fl/fl^;Sox1^Cre/+^* mutants (26.87±5.12) compared to controls (9.20±4.30, P=0.0169), while *Sox9^fl/fl^;Nestin-Cre* mutants were not affected (15.93±6.60, Turkey’s multiple comparison test, ANOVA P=0.0197). **(H-J)** Granule neurons EdU birth-dating experiment. (**H**) Double staining for PROX1 and EdU on E18.5 control, *Sox9^fl/fl^;Sox1^Cre/+^* and *Sox9^fl/fl^;Nestin-Cre* mutant embryos. (**I**) EdU was injected at E16.5, which corresponds to the first stage when this cell type appears in the developing DG, and samples harvested at E18.5. (**J**) The proportion of PROX1+EdU+/PROX1+ newly formed granule neurons is unchanged between controls and both *Sox9* mutants, indicating that *Sox9* deletion is not affecting granule neuron differentiation. DNE: dentate neuroepithelium; CH: cortical hem; 1ry: primary matrix; 2ry: secondary matrix; 3ry: tertiary matrix. Scale bars represent 50µm in (Ai-ii); 100µm in (Aiii-iv), (D) and (H).

**Figure S.7.**
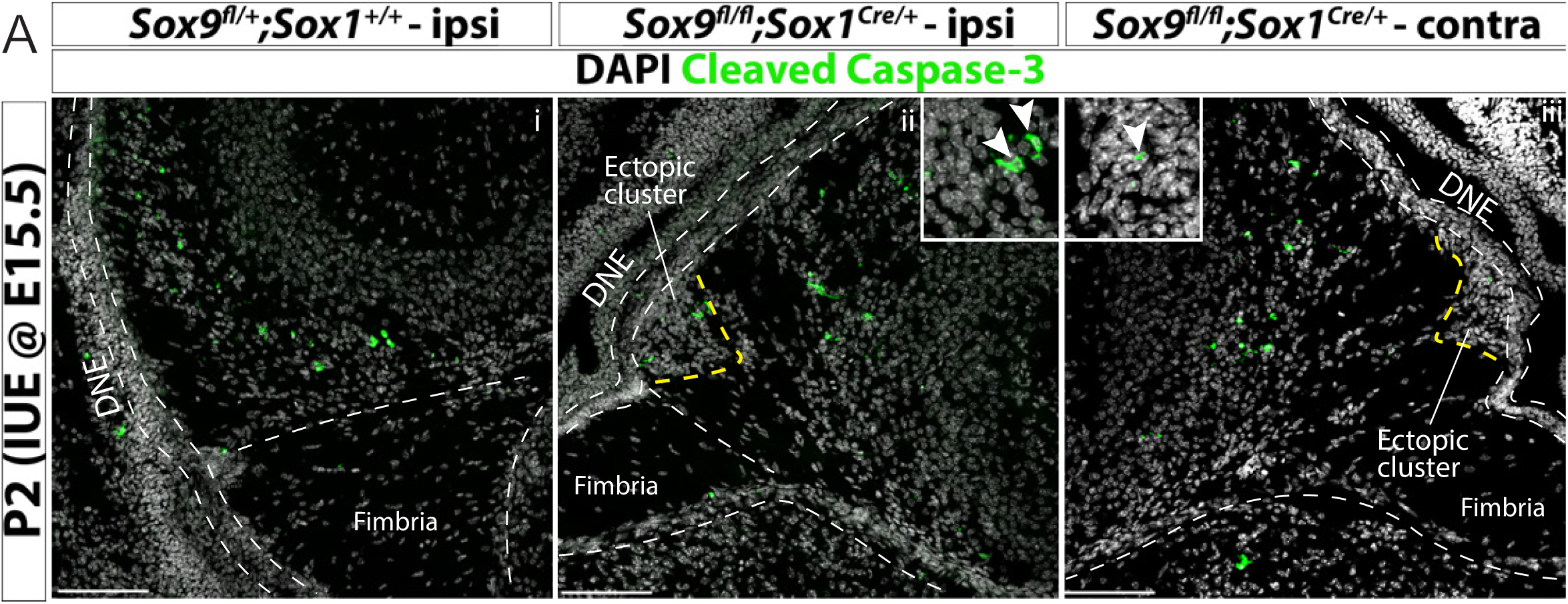
*In utero* electroporation does not compromise cell survival in the developing dentate gyrus of *Sox9* mutants. **(A)** Immunostaining for cleaved Caspase-3 in P2 *Sox9^fl/fl^;Sox1^Cre/+^* (ii,iii) pups and controls (i) after IUE at E15.5, comparing ipsi-(i,ii) and contra-lateral (iii) hemispheres to the injection, showing that cell death is not increased in *Sox9* mutants after IUE. Insets showing higher magnification of Cleaved Caspase-3 expressing cells in ectopic cluster. IUE: *in utero* electroporation; DNE: dentate neuroepithelium; ipsi: ipsilateral to injection; contra: contralateral to injection. Scale bars represent 100µm.

**Figure S.8.**
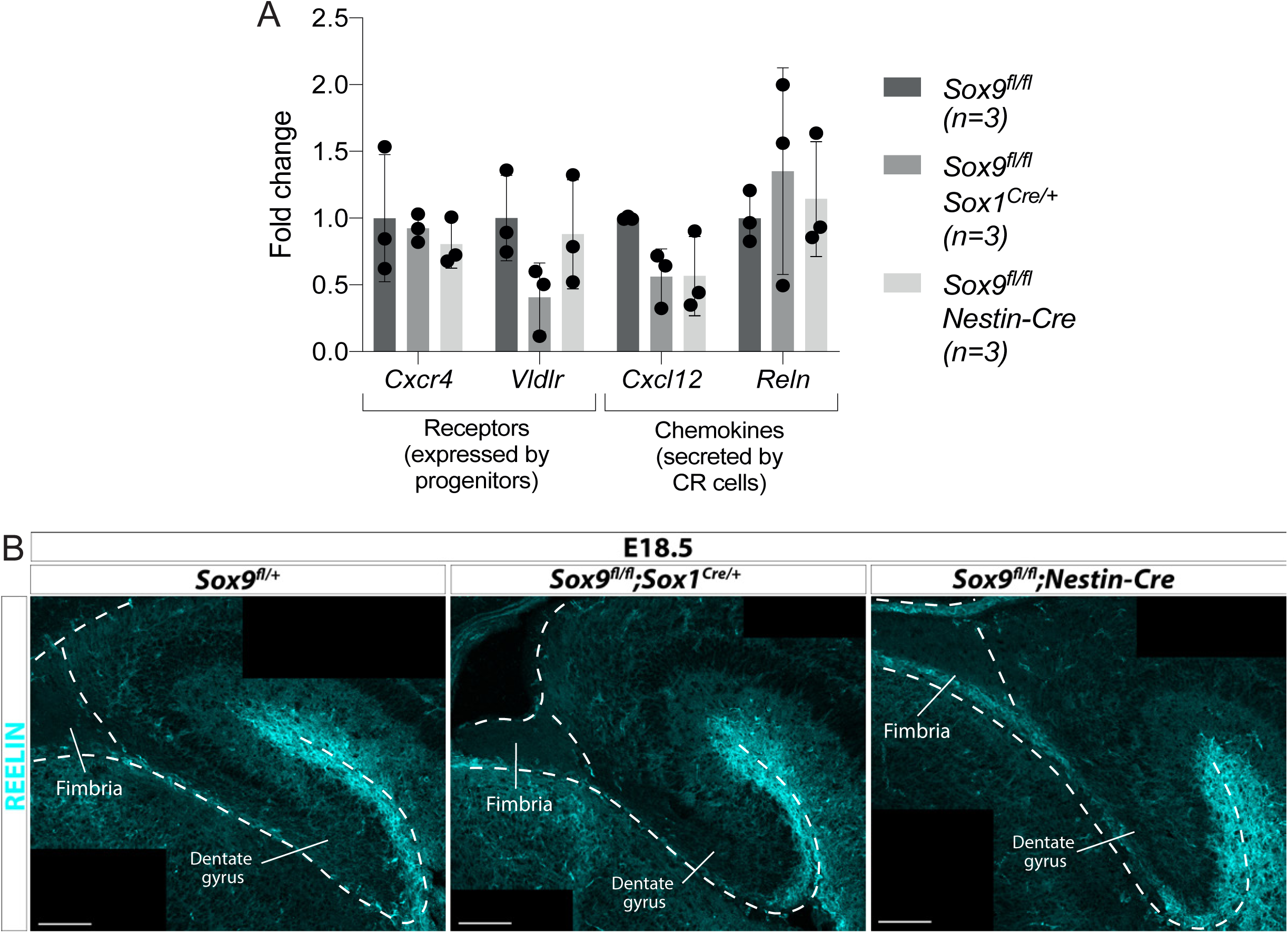
Migratory clues secreted by Cajal-Retzius cells and required during DG development are not affected in *Sox9* mutants. **(A)** Analysis by qPCR of the expression of secreted chemokines *Reln*, *Cxcl12* and their receptors *Vldlr*, *Cxcr4*. mRNA was extracted from E12.5 dissected archicortices of *Sox9^fl/fl^;Sox1^Cre/+^*, *Sox9^fl/fl^;Nestin-Cre* and control embryos. No significant difference was found in any of the genes analyzed in both *Sox9* mutants compared to controls. **(B)** Immunofluorescence for REELIN on E18.5 *Sox9^fl/fl^;Sox1^Cre/+^*, *Sox9^fl/fl^;Nestin-Cre* and control embryos. The expression pattern and intensity of REELIN staining appeared unaffected in both *Sox9* mutants compared to controls. Scale bars represent 100µm.

**Figure S.9.**
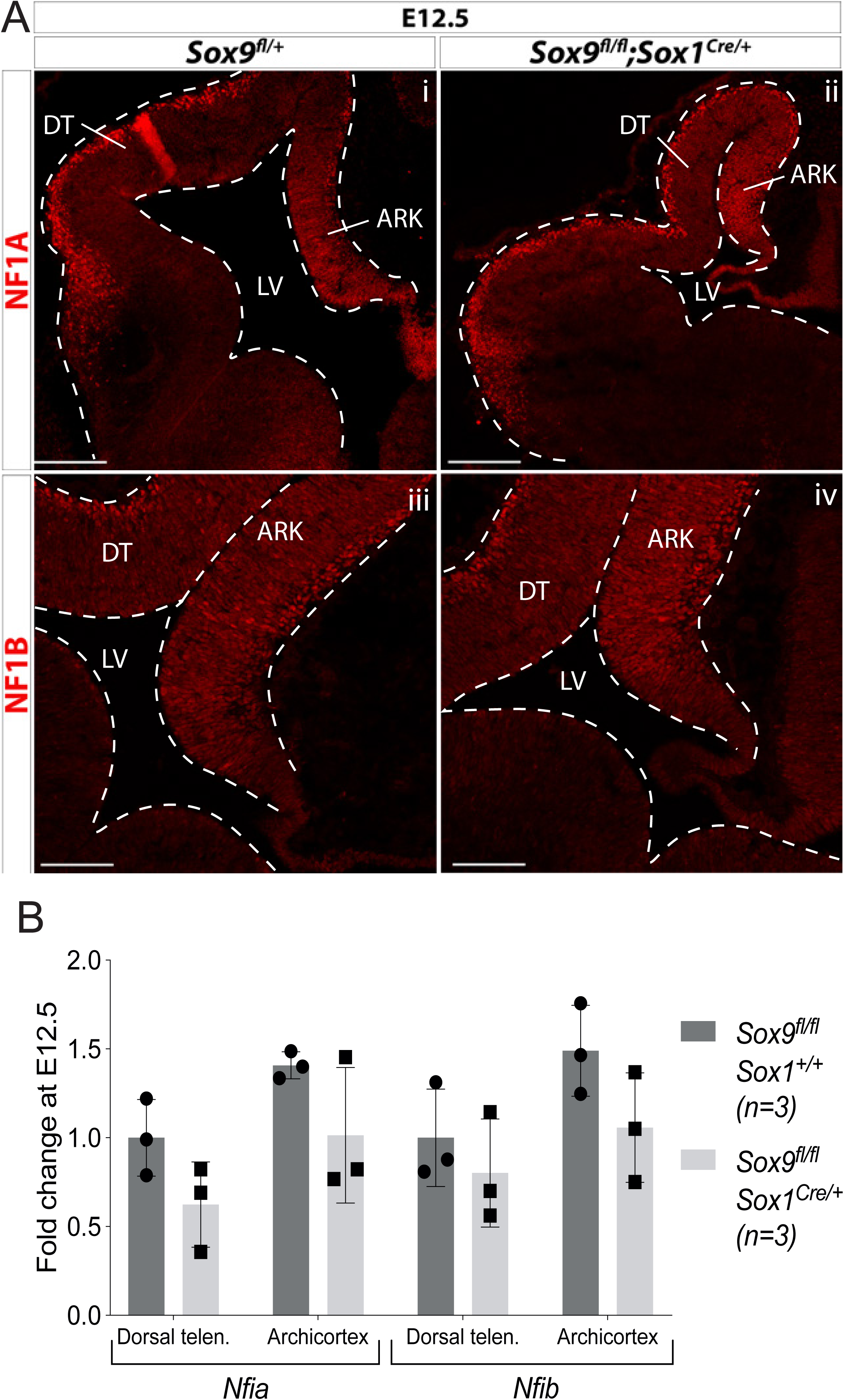
Early CNS-specific deletion of *Sox9* does not affect NF1A/B expression in the forebrain. **(A)** Immunofluorescence for NF1A (i,ii) or NF1B (iii,iv) on E12.5 *Sox9^fl/fl^*;*Sox1^Cre/+^* (ii, and control (i,iii) embryos. Levels of expression appear similar in both DT and ARK in *Sox9* mutants compared to controls. **(B)** Quantification of *Nfia/b* expression levels by qPCR on dissected E12.5 *Sox9^fl/fl^*;*Sox1^Cre/+^* and control dorsal telencephalons and archicortices. There is no significant difference in *Nfia/b* levels in *Sox9* mutants compared to controls. LV: lateral ventricle; DT: dorsal telencephalon; ARK: archicortex. Scale bar represent 200µm in (Ai-ii) and 100µm in (Aiii-iv).

**Figure S.10.**
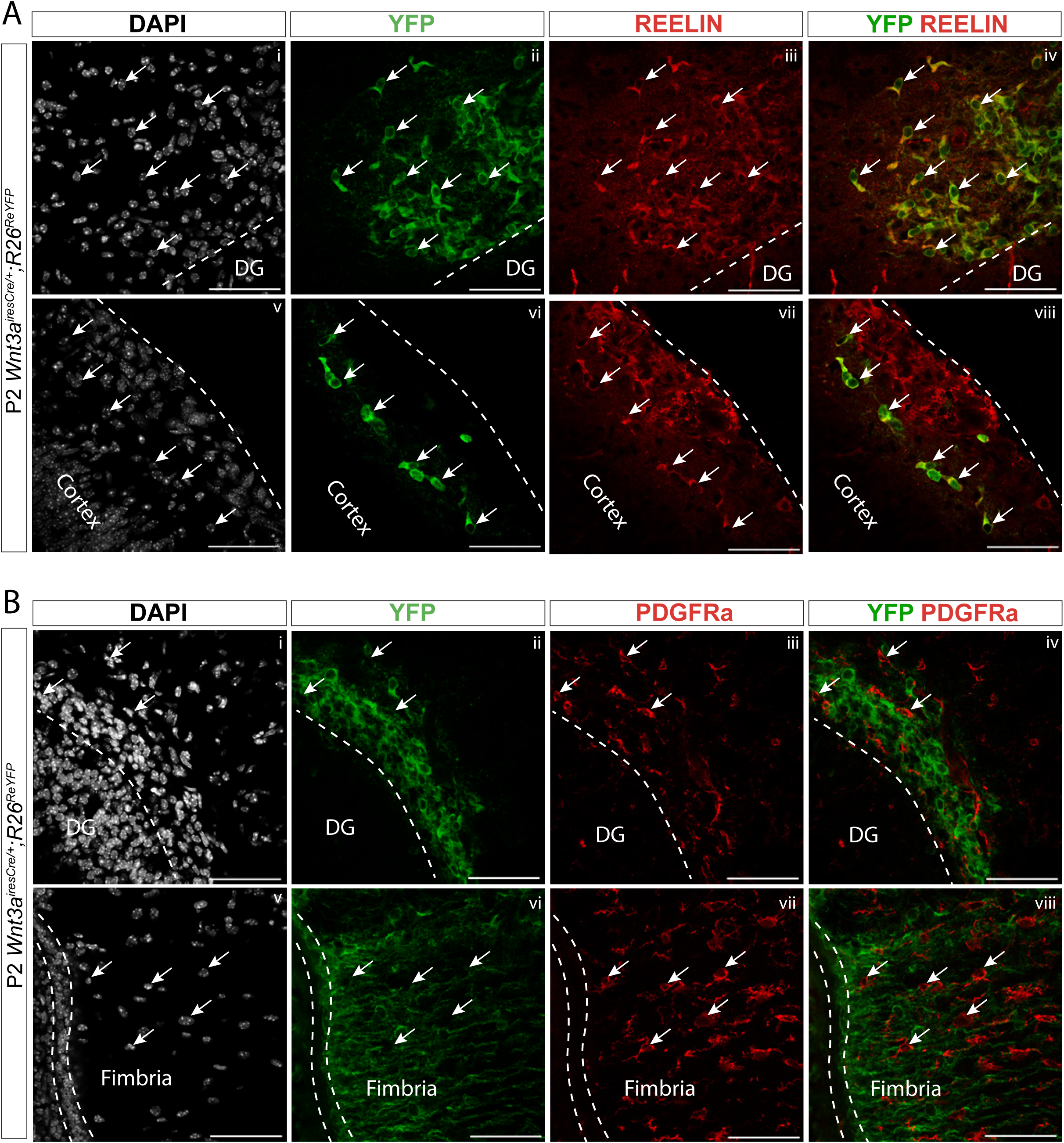
Lineage tracing analysis of CH-derived cells in *Wnt3a^iresCre/+^;R26^ReYFP^* pups. **(A,B)** Immunofluorescence for YFP and the Cajal-Retzius cell marker REELIN (**A**) or the oligodendrocyte precursor cells (OPCs) marker PDGFRa (**B**) on P2 *Wnt3a^iresCre/+^;R26^ReYFP^* brains. YFP+ cells express REELIN both around the DG (**A**i-iv) and in the outer layer of the cortex (**A**v-viii) (indicated by white arrows). Conversely, YFP is not expressed in PDGFRa+ cells, either around the DG (**B**i-iv) or within the fimbria (**B**v-viii) (indicated by white arrows). This suggests that Cajal-Retzius cells originate from the cortical hem, while OPCs do not. DG: dentate gyrus. Scale bar represent 50µm.

**Figure S.11.**
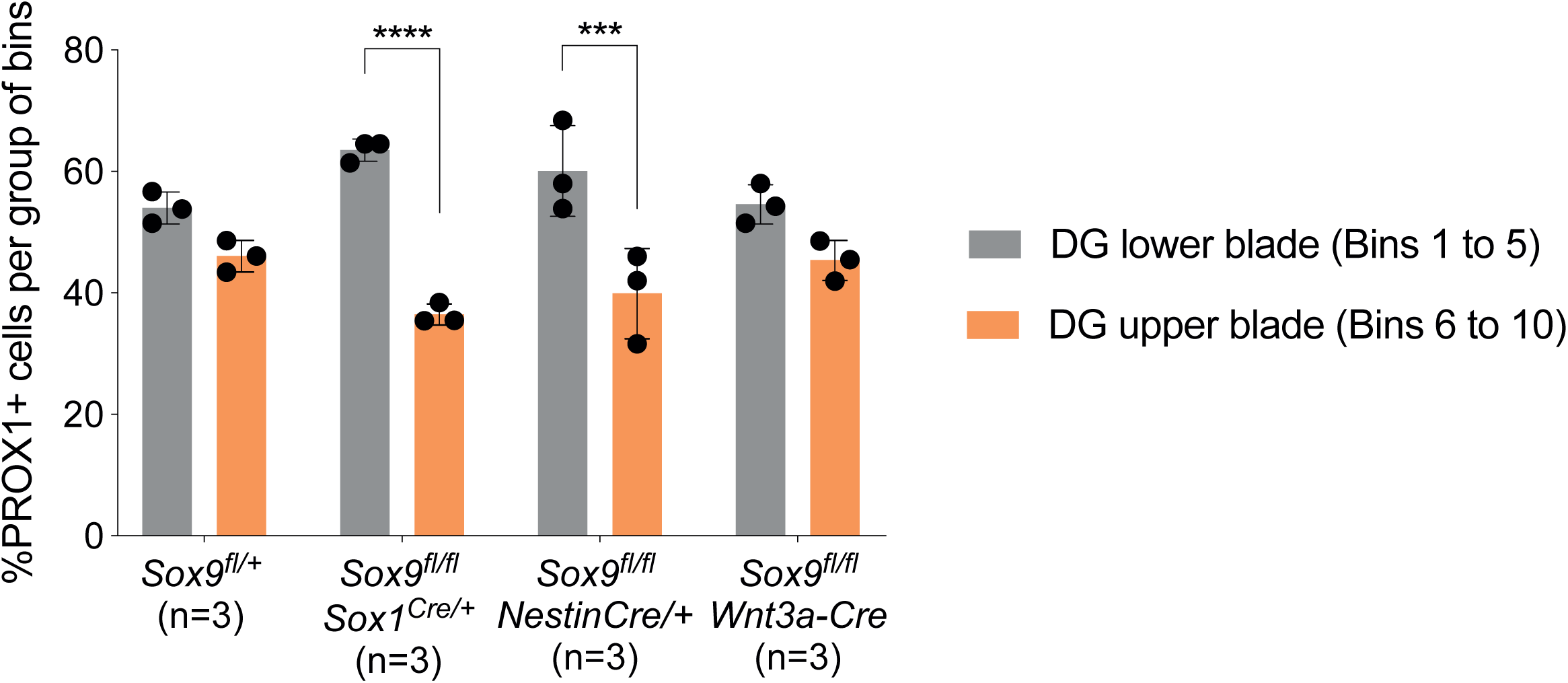
PROX1+ granule neurons upper/lower blade repartition in the 3ry matrix is not affected in *Sox9^fl/fl^;Wnt3a^iresCre/+^* mutants. Percentage of PROX1+ granule neurons positioned in the DG lower blade (bins 1 to 5), versus the DG upper blade (bins 6 to 10) in E18.5 controls, *Sox9^fl/fl^;Sox1^Cre/+^* and *Sox9^fl/fl^;Nestin-Cre* mutants (results from Fig. 2.N), and *Sox9^fl/fl^;Wnt3a^iresCre/+^* mutants (results from Fig. 7.C). In contrast with *Sox9^fl/fl^;Sox1^Cre/+^* and *Sox9^fl/fl^;Nestin-Cre* mutants, PROX1+ granule neurons are distributed normally in *Sox9^fl/fl^;Wnt3a^iresCre/+^* mutants.

